# Longitudinal tumor ecosystem mapping defines glioblastoma treatment trajectories

**DOI:** 10.64898/2026.03.04.709701

**Authors:** M Vanmechelen, P Nazari, J Beckervordersandforth, C Caprioli, DJG Leunissen, B Cole, C Bravo González-Blas, B Decraene, Y De Visser, G Shankar, M Verduin, D Pantano, S Bevers, T Moors, S Zielinski, J Telang, I Sebastian, J Messiaen, Y Van Herck, E Geens, DBP Eekers, A Claeys, M Derweduwe, Hausen A Zur, J Chui, FM Bosisio, F Weyns, T Daenekindt, C Oosterbos, P Van Eyken, M Govers, F Mennens, K Hovinga, S De Vleeschouwer, PM Clement, MPG Broen, M Vooijs, R Sciot, A Antoranz, J Pey, EJ Speel, A Hoeben, F De Smet

## Abstract

Glioblastoma remains an invariably recurring and lethal brain tumor shaped by complex interactions between malignant and microenvironmental cells. How these interactions evolve under first-line standard-of-care (SOC) therapy remains unclear. We performed spatial single-cell profiling of 671 paired newly diagnosed and recurrent glioblastoma samples from 96 patients to map tumor-ecosystem evolution and its clinical relevance. We delineated five distinct patient subgroups, each characterized by unique ecosystem trajectories that correlated with clinical outcome. Patients whose tumors transitioned into oligodendrocyte-progenitor-like niches with enhanced vascular integrity, oxygenation and an IL10/CCL/MHCII immunomodulatory environment experienced improved clinical prognosis. In contrast, progression towards hypoxic mesenchymal/astrocyte-like niches dominated by strong LGALS1/SPP1/TGFß immunosuppression predicted poor outcome. Furthermore, a subgroup of early-relapsing patients on SOC therapy that was characterized by a depletion of an antigen-presenting myeloid-cell-rich niche, exhibited diminished responsiveness to second-line lomustine therapy. Overall, mapping evolutions in glioblastoma ecosystems offers a novel framework for prognostic stratification and therapeutic guidance.

## Introduction

Adult glioblastoma (IDH^WT^ astrocytoma grade 4; GBM) are highly malignant, intrinsically resistant, and inevitably recurring brain tumors with dismal prognosis of merely 15 months despite intensive treatment^1,2^. First-line standard-of-care (SOC) treatment consists of a combination of maximal safe surgical resection followed by radiotherapy (RT) combined with alkylating chemotherapy (temozolomide, TMZ) ^1,2^. Methylguanine methyltransferase (*MGMT*) promoter methylation is the only well-established prognostic biomarker with predictive value for response to TMZ-based therapies^3^. When GBM tumors progress, lomustine chemotherapy is commonly used as second line treatment, although its effect on overall survival remains limited^4^. In addition, there is a lack of clear predictive biomarkers for lomustine response^5^.

The aggressiveness and lack of effective treatment strategies in GBM can be attributed to the highly heterogeneous and plastic nature of GBM, which easily confer resistance to both SOC and experimental treatments, for which clinical trials have failed over the past 20 years^6–8^. This heterogeneity exists across patients but also within the same tumor at multiple levels. At the tumor cell level, mixtures of various (epi)genomic, transcriptomic and proteomic cell states are typically observed, which are moreover highly plastic and dynamic enabling them to adapt to most therapeutic perturbations^9,10^. In addition, the tumor microenvironment (TME), which consists of resident and infiltrating immune cells, blood vessels, resident glial cells, neurons and extracellular matrix surrounding the tumor cells, shows extended heterogeneity and complexity^11,12^. Within the GBM ecosystem, microglia and tumor-associated macrophages (TAMs) are the most prevalent immune cells, which have been shown to fuel a strong pro-tumoral, immunosuppressive environment^13^, while T cells are comparatively scarce, often functionally exhausted, and largely excluded from the tumor core^14^.

To unravel insights into the resistance mechanisms of GBM to SOC therapy, studies have focused on brain tumor samples from baseline (*i.e.* newly diagnosed, treatment naive surgical samples; ND), or recurrent (REC) stages, and, where possible on longitudinal samples from the same patients^15–17^. Phenotypic shifts of tumor cells towards a mesenchymal (MES) phenotype were typically observed in cases with a worse prognosis^9,18–21^, likely driven by treatment induced clonal selection and adaptation to the local/immune microenvironment (e.g. reactive or hypoxic conditions)^22,23^. In addition, the spatial organization of the various cell types has also been shown to be relevant^12,24,25^. Novel technologies for spatially resolved single-cell mapping (e.g. spatial transcriptomics, multiplexed immunohistochemistry, and their combination) enable to profile tens to hundreds of biological markers in the spatial context of a single tissue section,^26,27^ allowing to study cell-cell interactions, define cellular ecosystems, and describe detailed aberrations in tissue architecture. Despite this progress, it remains unknown how classical, tumor-targeted therapeutics influence the interactions between tumors and their microenvironment, and how this contributes to responsiveness/resistance.

To map the variation of tumor-immune interactions in GBM upon SOC treatment, we have spatially characterized 2 multicentric cohorts of paired newly diagnosed (i.e. treatment-naïve) and post-treatment, recurrent tumor samples derived from 96 patients. This high-dimensional multiomics approach allowed us to quantitatively assess patient-specific trajectories of therapy-induced evolution in tumor architecture and ecosystem composition, enabling us to unravel the underlying molecular drivers while enabling the stratification of patients into novel clinically relevant subgroups. Finally, the observed trajectories in ecosystem composition also defined the responsiveness to lomustine as salvage therapy.

## Results

### Tissue architecture mapping reveals a change in tumor cell density at recurrence with increased immune interactions

To map changes in the cellular ecosystems of glioblastoma tumors, we collected a representative multicentric cohort of 52 GBM patients treated with SOC therapy, from which we collected paired tumor tissue samples (FFPE) at initial new diagnosis (ND) and recurrence (REC) and detailed longitudinal clinical, radiological and treatment information (Figure 1a,b, Table S1a,b, Note S1). *MGMT* promoter methylation and genomic profiling (n = 41 patients) highlighted the expected aberrations^28^, which could either be retained, appear, or disappear at recurrence in a patient-specific pattern (Figure 1c). By performing multiplexed immunohistochemistry (mIHC)^13,29,30^ with 38 relevant markers (Table S2), we mapped the longitudinal evolution of the spatial single-cell architecture in each GBM patient, generating a spatial single-cell proteomics GBM atlas of 4,788,453 cells across 357 tissue sections, provided with clinical annotation (Figure 1d,e, Figure S1-S2, Table S2-S3).

**Figure 1:**
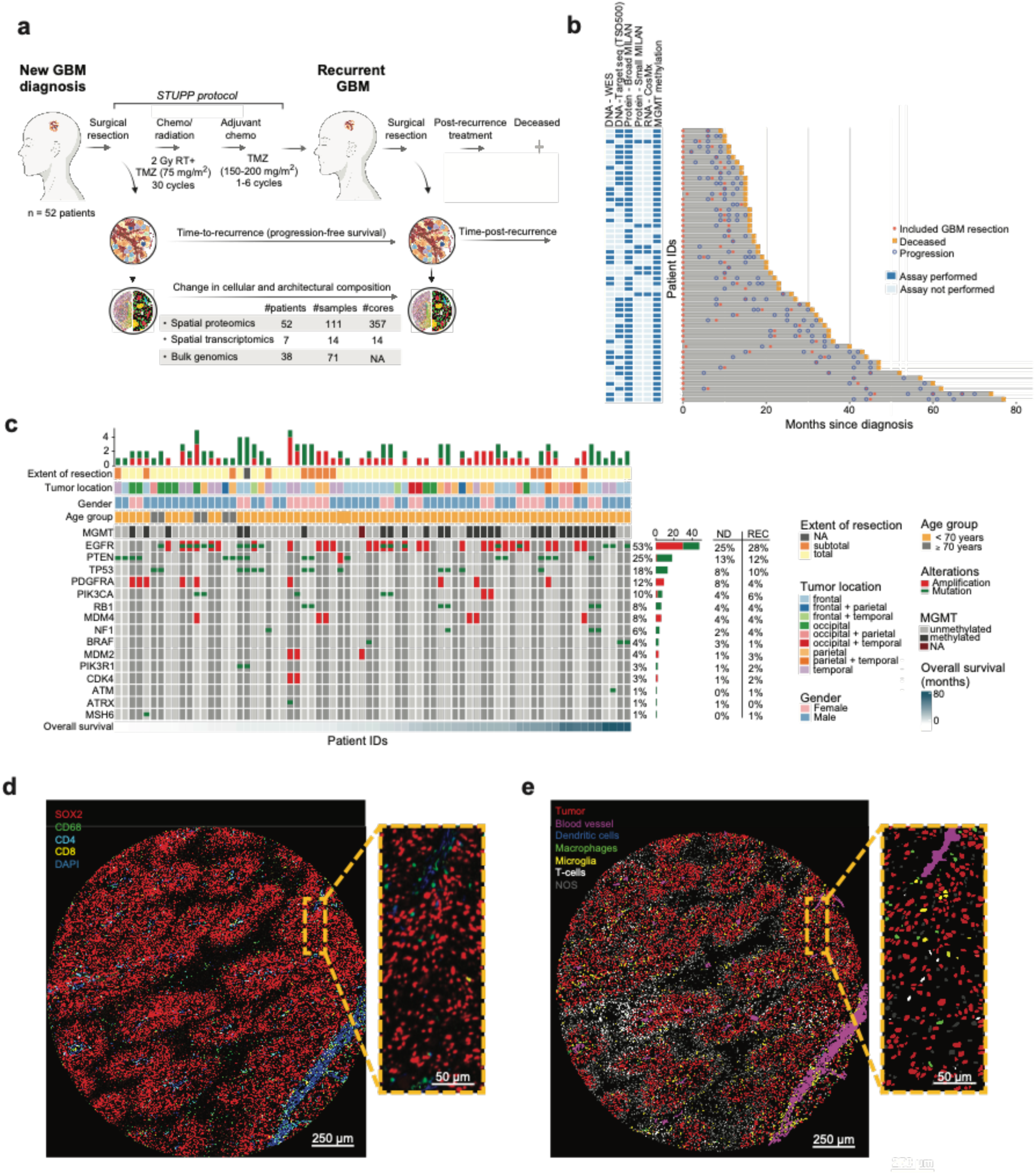
Overview of patient cohort and experimental approach. **a** Clinical course of glioblastoma patients from diagnosis through standard-of-care treatment (Stupp regimen). Overall, 52 patients were included in the discovery cohort, for which tumor samples from at least 2 time points (diagnosis and recurrency) have been profiled by spatial single cell proteomics (multiplex IHC, MILAN) and/or spatial transcriptomics. GBM, glioblastoma; RT, radiotherapy; TMZ, temozolomide; NA, not available. Image generated with Biorender.com. **b** Swimmer plot showing an overview of the analyzed patients, samples and tissue cores. The x-axis indicates the overall survival in months, while the y-axis highlights patients ranked from shortest to longest overall survival. For every patient, the different assays that have been performed are indicated. WES, whole exome sequencing; MILAN, Multiple iterative labeling by antibody neodeposition); MGMT, O-6-methylguanine-DNA methyltransferase. **c** Oncoprint summarizing the genetic background of patients with available information. The type and frequency of gene alterations and mutations are shown, combined with clinical characteristics such as extent of resection, location of the tumor, gender and age. MGMT, O-6-methylguanine-DNA methyltransferase; ND, newly diagnosed; REC, recurrent; NA not available. **d** Representative tissue core showing an overlay of the following stained markers: SOX2 (red), CD68 (green), CD4 (cyan), CD8 (yellow), DAPI (blue). The dashed box highlights a region of interest at higher magnification. **e** Left: Digital reconstruction of the same tissue core as in D, where the main cell types are annotated based on a consensus clustering approach. The dashed box highlights the same region of interest at higher magnification. NOS, not otherwise specified.

We identified a spectrum of distinct morphological patterns in overall tissue architecture, cellular densities, and vascularity, together with areas of tissue micro-necroses (*i.e*. small regions of necrosis encircled by pseudopalisading hypercellular neoplastic cells) (Figure 2a,b, S3a). Comparative analysis showed that ND specimens were typically composed of a higher proportion of high-density tumor regions compared to the REC resections (*P* = 9.56 x 10^-6^, Wilcoxon signed-rank test), in which low tumoral density areas were instead more abundant (*P* = 3.58 x 10^-2^, Wilcoxon signed-rank test) (Figure 2c,d). Accordingly, in the REC samples, the non-tumoral areas (typically composed of large clusters of immune cells that do not directly interact with tumor cells) became more abundant (*P* = 2.45 x 10^-4^, Wilcoxon signed-rank test) (Figure 2c,d). This was further confirmed upon investigating variations in the overall cell type composition, as at recurrence tumor cells became less abundant (*P* = 4.09 x 10^-3^, Wilcoxon signed-rank test) while macrophages and cytotoxic T cells, even though only present in low amounts, were instead relatively more enriched (*P* = 4.66 x 10^-3^, and *P* = 9.16 x 10^-4^ respectively, Wilcoxon signed-rank test) across the analyzed tissue areas (Figure 2e).

**Figure 2:**
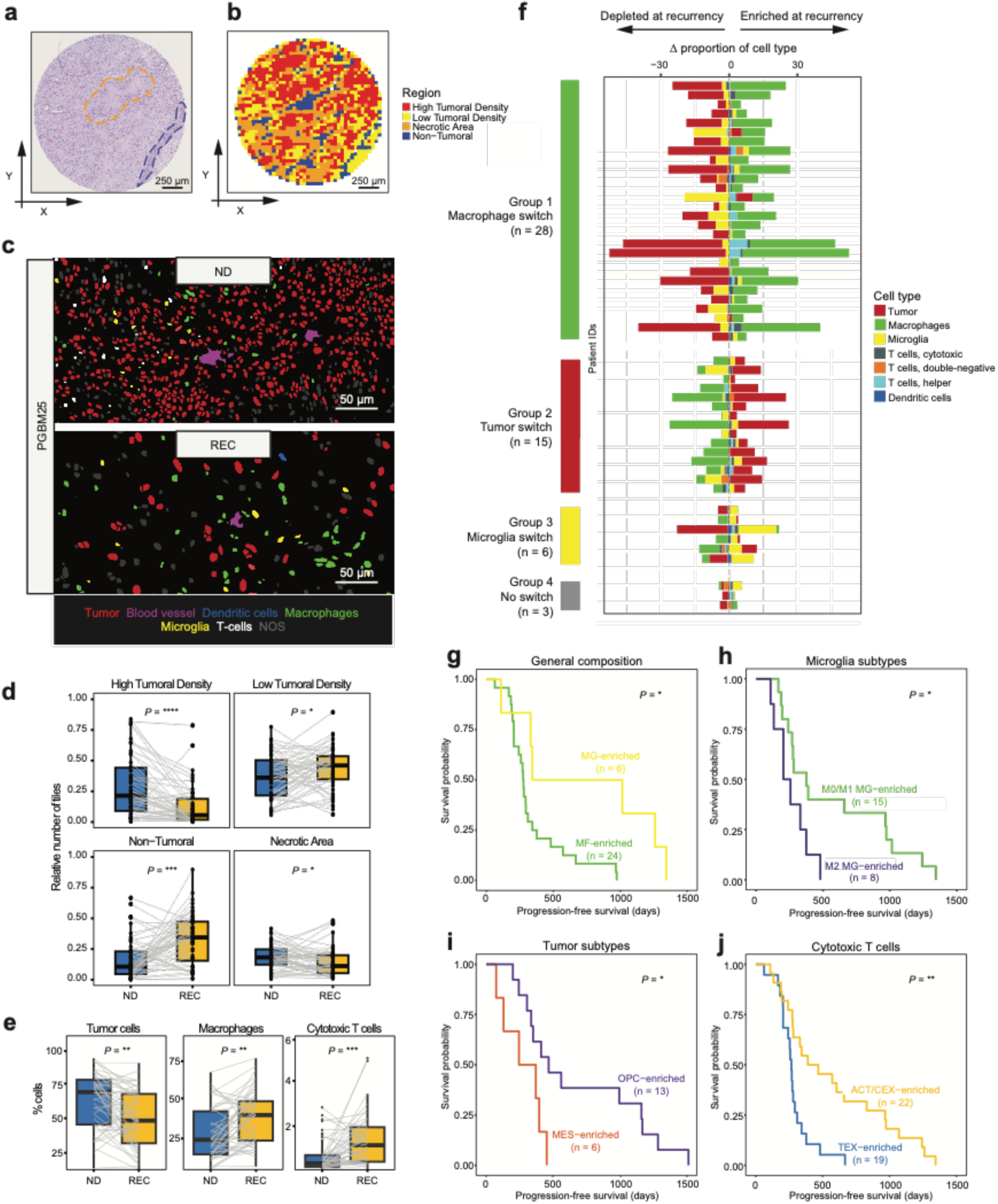
Patterns of ND-to-REC evolution in cell type composition and correlation to progression-free survival. **a** Histological image of a representative tissue core (same as in Figure 1d**,e**) by hematoxylin-eosin staining. The dashed lines highlight an area of pseudopalisading necrosis (orange) and a blood vessel (blue).**b** Density region metric classifying the GBM tumor architecture in 4 different regions based on the number of tumor cells in a square micro-region of 70 x 70 micron (same representative tumor core as in **a**). The defined regions are respectively ‘High tumoral density’ (tumor cells > 8), ‘Low tumoral density’ (0 < tumor cells ≤ 8), ‘Necrotic area’ (NOS cells > 8) and ‘Non-tumoral area’ (no tumor cells). **c** Example of ND-to-REC shift in cellular density and composition from a representative GBM patient, based on digital reconstruction from mIHC. Top: subregion of a newly diagnosed tissue specimen, showing high cellular density and prevalence of tumor cells. Bottom: subregion of recurrent tissue specimen from the same GBM patient, highlighting lower cellular density, depletion of tumor cells and higher prevalence of tumor-associated macrophages. ND: newly diagnosed, REC: recurrent; NOS, not otherwise specified. **d** Comparison of the distribution of the 4 different density regions (High tumoral density, Low tumoral density, Non-tumoral, Necrotic area) between newly diagnosed and recurrent tumor samples (evaluable on 46, 52, 52 and 51 patients respectively). Boxplots span from the first to third quartiles, middle line represents median. Significance was calculated by Wilcoxon signed-rank test for paired samples. ND: newly diagnosed, REC: recurrent. ns, *P* > 0.05; *, *P* ≤ 0.05; **, *P* ≤ 0.01; ***, *P* ≤ 0.001; ****, *P* ≤ 0.0001. **e** Comparison of proportions for main cell types between newly diagnosed and recurrent tumor samples (47 patients); only significant tests are shown. Boxplots span from the first to third quartiles, middle line represents median. Significance was calculated by Wilcoxon signed-rank test for paired samples. ND: newly diagnosed, REC: recurrent. ns, *P* > 0.05; *, *P* ≤ 0.05; **, *P* ≤ 0.01; ***, *P* ≤ 0.001; ****, *P* ≤ 0.0001. **f** Patterns of evolution in cell type composition between newly diagnosed (ND) and recurrent (REC) tumor samples for each individual GBM patient. The delta value (Δ, on the x-axis) shows the ND-to-REC difference in the proportion of each cell type, which allows grouping patients with similar shifts into different evolutionary patterns. **g-j** Kaplan-Meier plots showing differences between patient groups as stratified by ND-to-REC shifts in general cellular composition (**g**), tumoral subtypes (**h**), microglia subtypes (**i**), and cytotoxic T cells (**j**). MF, macrophages; MG, microglia; OPC, oligodendrocyte precursor-like; MES, mesenchymal-like; ACT, activated T cells ; CEX, chronic exhausted T cells; TEX, terminally exhausted T cells. Significance was calculated by Log-Rank test. ns, *P* > 0.05; *, *P* ≤ 0.05; **, *P* ≤ 0.01; ***, *P* ≤ 0.001; ****, *P* ≤ 0.0001.

### Longitudinal evolution of cellular composition patterns correlates to progression-free survival

Next, we classified all cells into specific functional (sub)states, focusing on tumoral (including molecular substates such as astrocyte-like (AC), oligodendrocyte-precursor like (OPC), neuronal-precursor like (NPC) and mesenchymal (MES)-like cells, and other general programs such as stemness and proliferation) and immune cell states [such as activation/exhaustion of T cells, polarization of macrophages (MF) / microglia (MG)] (Figure S1c-j). Interestingly, withing the AC-like population, a large subgroup of cells was also EGFR+, which we investigated separately (AC-like EGFR+ (ACE)), in line with previous work^31^.

The analysis of longitudinal changes in cellular composition between diagnosis and recurrence allowed us to classify patients based on similar patterns of evolution after first line treatment (Figure 2f and S3b-f). Comparison of progression-free survival (PFS) (*i.e.* the time to recurrence after initial resection) between these groups revealed that an increase in relative abundance of MG cells as opposed to an enrichment of MF was associated with delayed recurrence [median survival 678 days (CI95% 330-NA) vs 274 days (CI95% 241-342), *P* = 1.5 x 10^-2^, Log-Rank test] (Figure 2g). Grouping based on functional polarization within the MG compartment allowed to further stratify patients exhibiting slow or fast recurrence (increase in M0/M1-like MG or M2-like MGs at progression or, respectively) [median survival 377 days (CI95% 269-1015) vs 230 days (CI95% 204-NA), *P* = 3.5 x 10^-2^, Log-Rank test] (Figure 2h). Moreover, patients showing enrichment in OPC-like tumor cells after SOC treatment showed longer PFS, whereas increased MES-like prevalence correlated with shorter PFS [median survival 389 days (CI95% 280-NA) vs 256 days (CI95% 108-NA), *P* = 2.4 x 10^-2^, Log-Rank test] (Figure 2i). Finally, even though the abundance of CD8⁺ T cells was low compared to many other cancer types^32^, tumors evolving towards a relatively higher proportion of activated/chronically exhausted cytotoxic T cells (CD8Act/CD8Cex) were associated with prolonged times to recurrence compared to those enriched in terminally exhausted T cells (CD8Tex) [median survival 428 days (CI95% 280-967) vs 266 days (CI95% 242-342), *P* = 1.8 x 10^-3^, Log-Rank test] (Figure 2j). Importantly, these changes were most apparent in the low-density tumor areas where most interactions between tumor-immune cells are ongoing, highlighting the need for spatial profiling.

### Integrative cellular neighborhood analysis identifies clinically relevant cellular niches

To more carefully investigate tumor architecture, we next identified 45 cellular neighborhoods (CNs) across the entire cohort, each with a specific composition of tumor and immune cells (Figure 3a-b, S4a) that aligned with the observed staining patterns and distribution across the high/low-density tumoral, non-tumoral and necrotic areas (Figure S4b). We found that the proportion of tumor cells contributing to each community widely varied (from ∼10% in CN7 to ∼95% in CN15), AC-like tumor cells being the most represented subtype (Figure 3a, S4a).

**Figure 3:**
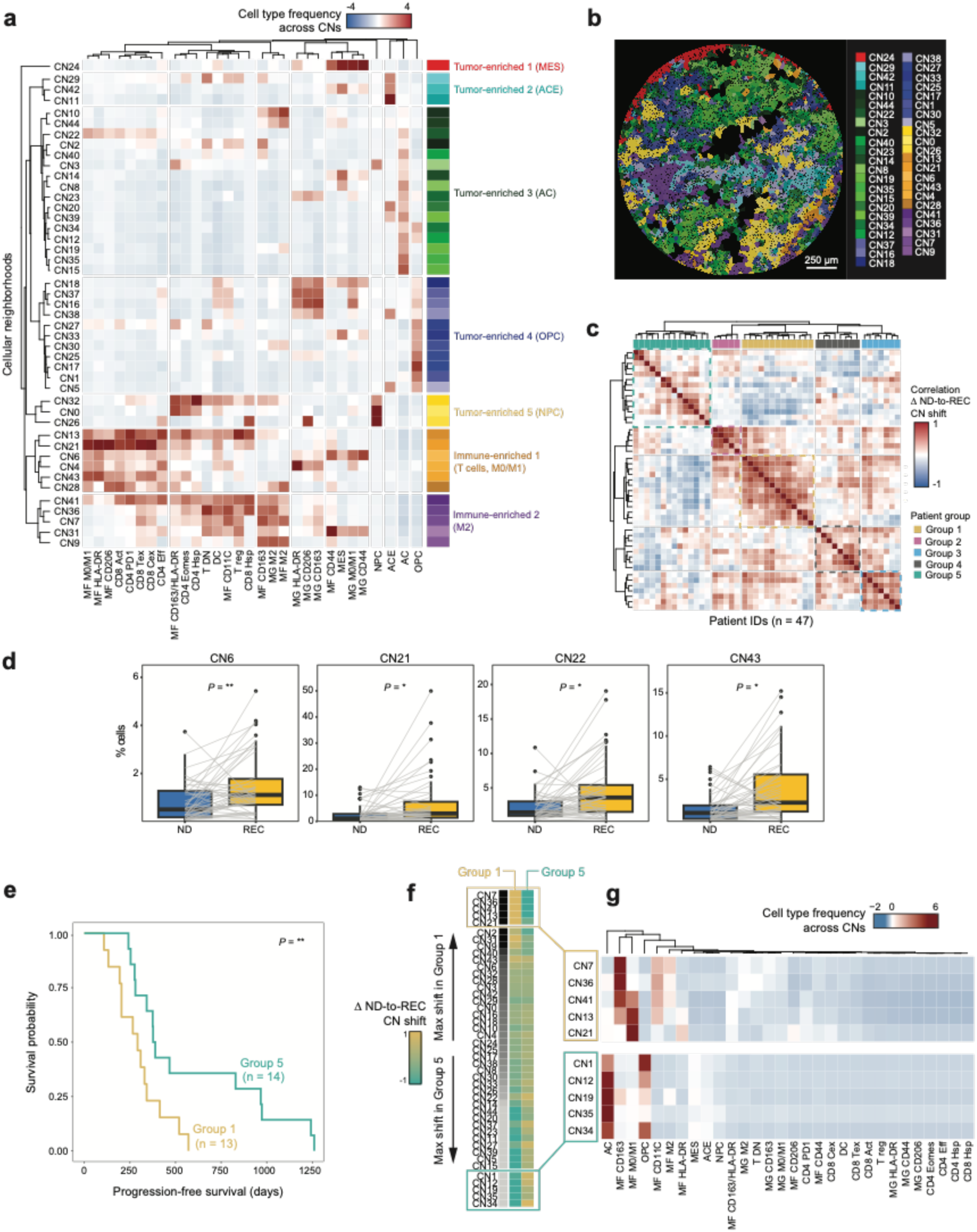
Patterns of ND-to-REC evolution in cellular neighborhood composition and correlation to progression-free survival. **a** Normalized frequency of cell types across the 45 cellular neighborhoods (CNs). Based on clustering, CNs are grouped in patterns of similarity in most enriched cell types, as denoted by the color code in the side bar. From top to bottom: red, tumor-enriched CNs, with MES-like dominance and CD44+ MF, CD44+ MG and M0/M1 MG; cyan, tumor-enriched CNs, with ACE-like dominance; green, tumor-enriched CNs, with AC-like dominance; blue, tumor-enriched CNs, with OPC-like dominance; yellow, tumor-enriched CNs, with NPC-like dominance; orange, immune-enriched with terminally exhausted CD8+ T cells, chronic exhausted CD8+ T cells, activated CD8+ T cells, PD1+ CD4 T cells, effector CD4+ T cells and atypically polarized macrophages (CD206+, M0/M1 and HLADR+); purple, immune-enriched with canonical CD163+ M2-like polarized macrophages and microglia cells. AC, astrocyte-like; ACE, astrocyte-like (+EGFR expression); CD4 eff, effector CD4+ T cells; CD4 PD1, PD1+ CD4 T cells; CD8 Act, activated CD8+ T cells; CD8 Tex, terminally exhausted CD8+ T cells; CD8 Cex, chronic exhausted CD8+ T cells; CNs, cell neighborhoods; DC, dendritic cells; MES, mesenchymal-like; MF, macrophages; MG, microglia; NPC, neural-progenitor-like; OPC: oligodendrocyte-progenitor-like; TCY, cytotoxic T cells; T DN, double-negative T cells; TH, helper T cells; T reg, regulatory T cells. **b** Spatial distribution of cellular neighborhoods (CNs) in a representative tissue core (same as in Figure 1d**,e** and Figure 2a**,b**). CNs are represented as Voronoi diagrams and colored based on the same scheme as in panel **a**. **c** Patient stratification by Pearson correlation of the ND-to-REC shift in CN composition. CNs, cell neighborhoods; ND, newly diagnosed; REC, recurrent. **d** Boxplots of shift in CN proportions at diagnosis and recurrence for each patient (47 patients); only significant tests are shown. Significance of differences was tested by Wilcoxon signed-rank test for paired samples. ND: newly diagnosed, REC: recurrent, CNs: cell neighborhoods. *, *P* ≤ 0.05; **, *P* ≤ 0.01. **e** Kaplan Meier plot showing difference in progression-free survival for patients belonging to group 1 and 5 according to the stratification performed in **c**. Significance was estimated by Log-Rank test. **, *P* ≤ 0.01. **f** ND-to-REC shift of each CN in group 1 and group 5. CNs are ranked based on the difference between group 1 and 5. ND: newly diagnosed, REC: recurrent, CNs: cell neighborhoods. **g** Normalized frequency of cell types across the top 5 most differential CNs between group 1 and group 5. ND: newly diagnosed, REC: recurrent, CNs: cell neighborhoods.

Next, we investigated the colocalization patterns of specific cell types across both tumor-enriched and immune-enriched CNs (Figure 3a, S4a,c). MES-like tumor cells colocalized primarily with CD44+ MF and CD44+ MG, while other tumoral subtypes (*i.e.* AC, ACE, NPC and OPC-like) showed more diverse interactions in the context of both immune-enriched or deprived CNs (Figure 3a, S4a,c). We found colocalization of the canonical CD163+ M2-like polarized MFs and MGs, in addition to two more complex patterns of immune-enriched CNs: pattern 1, composed of T cells (Terminal exhausted CD8+ T cells (CD8Tex), chronic exhausted CD8+ T cells (CD8Cex), activated CD8+ T cells (CD8Act), PD1+ CD4 T cells (CD4PD1) and effector CD4+ T cells (CD4eff) cells) and atypically polarized MFs (CD206+, M0/M1 and HLADR+ MFs), and pattern 2 composed of CD11c+ MFs, CD163+/HLADR+ MFs, and more rare cell types including Tregs, HSP70+ and EOMES+ CD4 T cells and dendritic cells (DCs) (Figure 3a, S4a,c). In CNs where activated CD8⁺ T cells were present, we generally observed a chronically or terminally exhausted phenotype (Figure S4d). Interestingly, we found that some immune pattern 1 communities (*i.e.* cytotoxic T cells and M0/M1 polarized myeloid cells) consistently showed relative increase at recurrence, namely CN43 and CN21 (both purely immune-enriched), CN6 (colocalizing with MES-like tumor cells), and CN22 (colocalizing with AC-like tumor cells) (Figure 3d).

To further investigate the impact of tissue structure evolution, we stratified patients with available survival data based on their longitudinal changes in neighborhood-based composition and identified five subgroups (Figure 3c). Interestingly, these groups showed a gradual shift in PFS (Figure S4e), the maximum difference being between groups 1 and 5 [(median PFS 292 days (CI95% 204-NA) vs 383 days (CI95% 342-1242) respectively, *P* = 1.91 x 10^-2^, Log-Rank test)] (Figure 3e, Table S4). Here, group 1 was characterized by an enrichment in non-tumoral immune neighborhoods (CN7, CD36, CN21, CN41 and CN13), which all contain high amounts of M2 polarized MFs and MGs (Figure 3f,g), in addition to a relative enrichment in AC-like or MES-like tumor cells. Conversely, patients in group 5 showed enrichment for OPC/AC-driven communities with limited macrophage polarization, in addition to HLA-DR positive macrophages (alone or in combination with CD163) (Figure 3f,g).

When integrating genomic features, we observed a trend toward a significant association only for EGFR amplification status (Fisher’s exact test, *P* = 0.09), driven primarily by an increased incidence of wild-type to amplified EGFR transitions in group 5, which were absent in the other groups (Table S5).

### Spatial multiome profiling identifies functional tumor-driven niches

To achieve a deeper characterization of the GBM cellular niches, we combined spatial transcriptomics and spatial proteomics on consecutive FFPE sections of 14 paired tissue samples from 7 GBM patients, in which we measured the expression of a panel of 1,008 genes (Table S6) in addition to 24 protein markers (Table S3), and integrated both modalities (Figure 4a, S5a-d,g, Table S7). Specifically, we were interested in understanding whether the different tumoral niches might exhibit functional polarization, based on the importance of secreted factors (such as growth factors, cytokines and chemokines) in modulating both tumor and immune cells (Figure 4, S5e,f,h,i). We found that tumor-derived C-C Motif Chemokine Ligand 2 (*CCL2)* was preferentially expressed in areas enriched with MES-, AC- and ACE-like tumor cells, while the OPC-like and the NPC-like niches were enriched for tumor-derived Natriuretic peptide C (*NPPC*) (Figure 4b). Importantly, such cytokine-driven polarization pattern was consistent across the integrated dataset combining proteomics and transcriptomics (Figure 4b and S5h,i) but also transcriptomics alone (Figure 4c,d). In addition, single-cell multiomics on the same tissue section (see *Methods*) further confirmed that *NPPC* expression colocalized with OPC-like (OLIG2^high^) tumor cells, while *CCL2* expression was enriched in mesenchymal (CD44^high^) or AC-like tumor areas (Figure 4f,g). We also found that *NDRG1*, a marker for hypoxia^33^, was enriched close by the *CCL2+* areas, while more limited in the *NPPC+* areas (Figure 4f,g).

**Figure 4:**
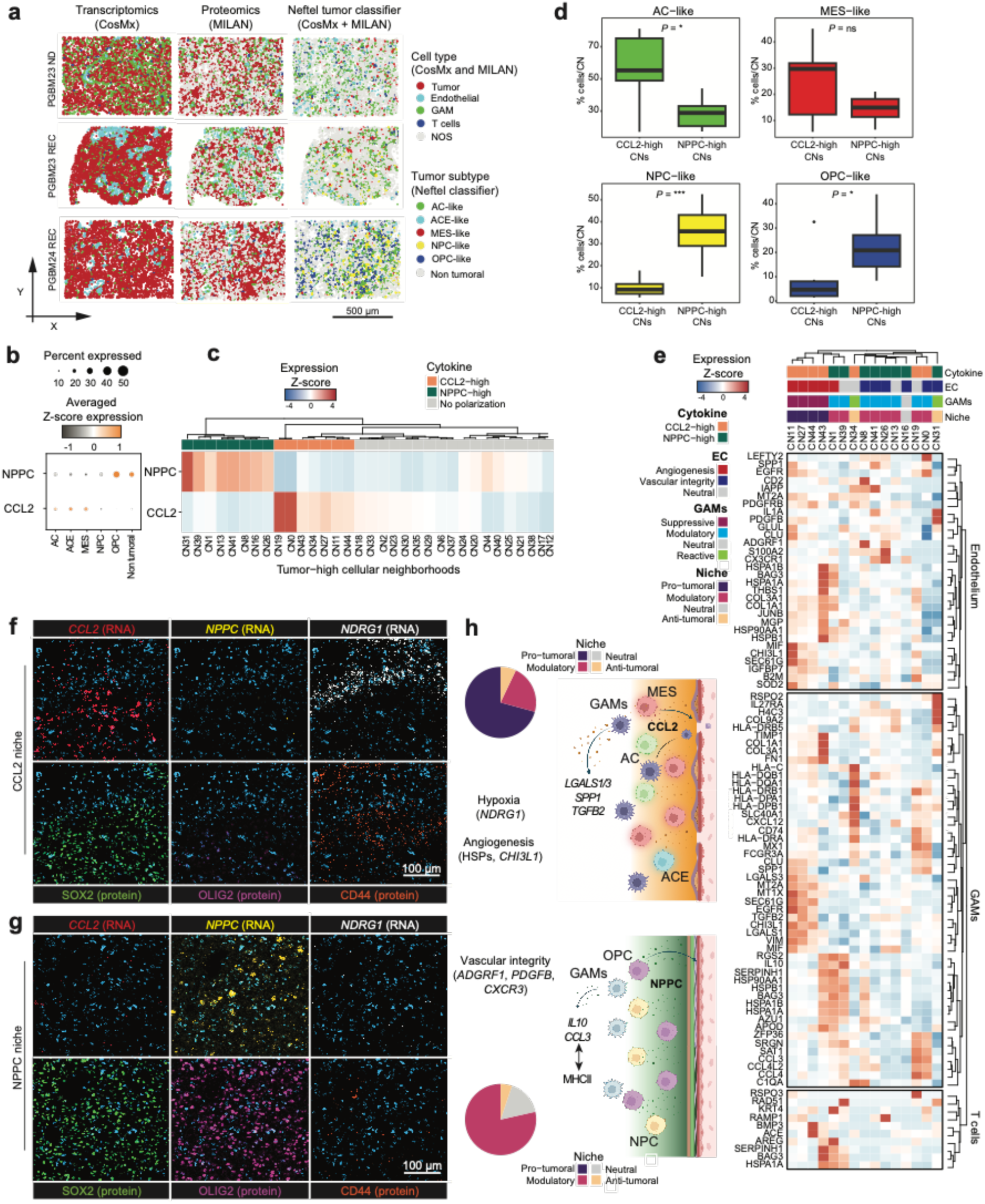
Mapping niche-specific cytokine networks. **a** Integration of single-cell spatial transcriptomics and proteomics on consecutive tissue slides. For the tumoral compartment, Neftel-like areas/tiles were defined using spatial proteomics data, dominated by one of the different tumoral subtypes from the single cells analysis. A subset of 3 highly correlating tissue sections is shown (see also Figure S5g). AC: astrocyte-like, ACE: astrocyte-like (+EGFR expression), GAMs: glioblastoma-associated macrophages, MES: mesenchymal-like, NOS: not otherwise specified, NPC: neural-progenitor-like, OPC: oligodendrocyte-progenitor-like. **b** Dotplot showing the average normalized expression of *CCL2* and *NPPC* in the integrated spatial transcriptomic dataset across AC, ACE, OPC, NPC and MES tumor cell populations, as well as non-tumoral populations. The size of the dots represents the percentage of cells expressing the marker, while the color indicates the intensity of expression. **c** Heatmap showing *CCL2-* and *NPPC*-polarization patterns across tumor-enriched CNs (independently scored from spatial transcriptomics data). CNs, cellular neighborhoods. **d** Boxplots showing the difference in proportion of *CCL2*-(n = 7) vs *NPPC*-polarized (n = 8) tumor-enriched CNs for each Neftel tumor subtype, from spatial transcriptomics data. AC: astrocyte-like, MES: mesenchymal-like, NPC: neural-progenitor-like, OPC: oligodendrocyte-progenitor-like. Significance of differences was tested by Wilcoxon rank-sum test. ns, *P* > 0.05; *, *P* ≤ 0.05; **, *P* ≤ 0.01; ***, *P* ≤ 0.001; ****, *P* ≤ 0.0001. **e** Heatmap showing, for each *CCL2*-vs *NPPC*-polarized tumor-enriched CN, the top 10 marker genes across the various tumor microenvironment components (from top to bottom, endothelial cells, GAMs, and T cells). CNs are labelled by their cytokine polarization, functional annotation of endothelial cells and GAMs, and overall pro-vs anti-tumor functionality of the cytokine niche (top annotation bars). CNs, cellular neighborhoods; GAMs, glioblastoma-associated macrophages; EC, endothelial cells. **f-g** Composite images of selected RNA and protein markers detected on the same GBM tissue section by RNAscope™ and seqIF™ profiling. Areas enriched with *CCL2* transcripts (**e**) also overexpress *NDRG1* (a hypoxia marker) and are preferentially populated by CD44^high^ (*i.e.,* MES-like) tumor cells, while areas enriched with *NPPC* transcripts (**f**) make a less hypoxic environment showing higher prevalence of OLIG2^high^ (*i.e.,* OPC-like) tumor cells. **h** Proposed functional model of CCL2-high vs NPPC-high niches, summarizing findings from **b** and **d-g**. The pie charts show the prevalence and overall pro-vs anti-tumor functionality of each CN within the CCL2-high and NPPC-high niche, respectively. The CCL2-high niche, enriched with MES- and AC-like tumor cells, consists of a hypoxic environment with prevailing pro-tumoral features, including dysregulated angiogenesis and immunesuppressive GAMs. Conversely, the NPPC-high niche exhibits less polarized features, with enhanced integrity of vascular structures, less hypoxia, and mixed GAMs phenotypes suggesting ongoing immune modulation. Image generated with Biorender.com.

Considering the known role of NPPC in maintaining vascular integrity^34^, we investigated whether *NPPC* vs *CCL2* cytokine polarization might impact on endothelial function. We found that endothelial cells in *NPPC+,* tumor-enriched cellular neighborhoods enriched for *ADGRF1* expression, a G-protein coupled receptor that enhances tight junction formation^35^, and *CXCR3* and *PDGFB,* both markers that ensure proper coverage by pericytes and the formation of a functional blood-brain barrier^36^, while in the *CCL2+* cellular neighborhoods the endothelium was characterized by a high stress response and CHI3L, both hallmarks of enhanced angiogenic activity^37^ (Figure 4e, Table S11). Altogether, this suggests that GBM tumor niches prime two different ecosystems: one dominated by a hypoxic environment enriched with MES/AC/ACE tumor cells producing high levels of *CCL2,* as opposed to an OPC/NPC-driven ecosystem with enhanced *NPPC*-induced vascular integrity and reduced hypoxia (Figure 4h). Spatial transcriptomics further identified that macrophages in *CCL2+* tumor-enriched communities showed transcriptional overexpression of genes related to M2 polarization and immunosuppression, like *TGFB2*^38^, *SPP1*^39^ and *LGALS1*^40^ (Figure 4e, Table S11). In the *NPPC+* environment, on the other hand, we identified macrophages with distinctive transcriptional signatures combining proteostasis support (*HSP90*, *HSP70*/*BAG3*), immunoregulatory signaling (*IL-10* or *IL27RA*), matrix-remodeling capacity (*SERPINH1*), complement mediation (*C1QA*), and controlled secretory/chemokine elements (*CCL3*, *CCL4*, *CCL4L*, among others) (Figure 4e). Far from being a purely immunosuppressive environment, the co-existence of *CCL3/4+* populations suggests that this ecosystem retains some capacity to modulate pro-inflammatory signals, in line with the various myeloid programs described in GBM^41^.

### Patterns of evolution in cellular neighborhood composition allow to stratify responsiveness to second line treatment with lomustine

Finally, we investigated whether the various patterns of evolution in tumor composition might also impact on responsiveness to post-recurrence therapy (Figure 5a, Table S12). For patients in this study cohort, second line treatment consisted of three possible strategies: (i) best supportive care (n = 10), (ii) lomustine (CCNU 110mg/m^2^) (n = 18) and (iii) rechallenge with a second course of temozolomide (150-200mg/m^2^) (n = 19). The latter was mostly administered to patients with initial good response to the Stupp regimen, while lomustine was given as salvage therapy to poor responders fit enough to receive further treatment [(median PFS 571 days (CI95% 389-1070) vs 271 days (CI95% 241-375) respectively, *P* = 4.7 x 10^-4^, Log-Rank test)] (Figure 5b). Also, when considering post-recurrence survival, patients receiving a second temozolomide challenge performed better than both those treated with lomustine or those only receiving supportive palliative care [(median survival 411 days (CI95% 369-825) vs 232 days (CI95% 174-454) and 220 days (CI95% 168-NA) respectively, *P* = 3.9 x 10^-2^, Log-Rank test)] (Figure 5c). Of note, patients undergoing temozolomide rechallenge showed a higher prevalence of *MGMT* promoter methylated tumors (66%, 31% and 25% of evaluable patients receiving temozolomide, lomustine and supportive care, respectively; *P* = 5.8 x 10^-2^, Fisher’s exact test) (Table S13).

**Figure 5:**
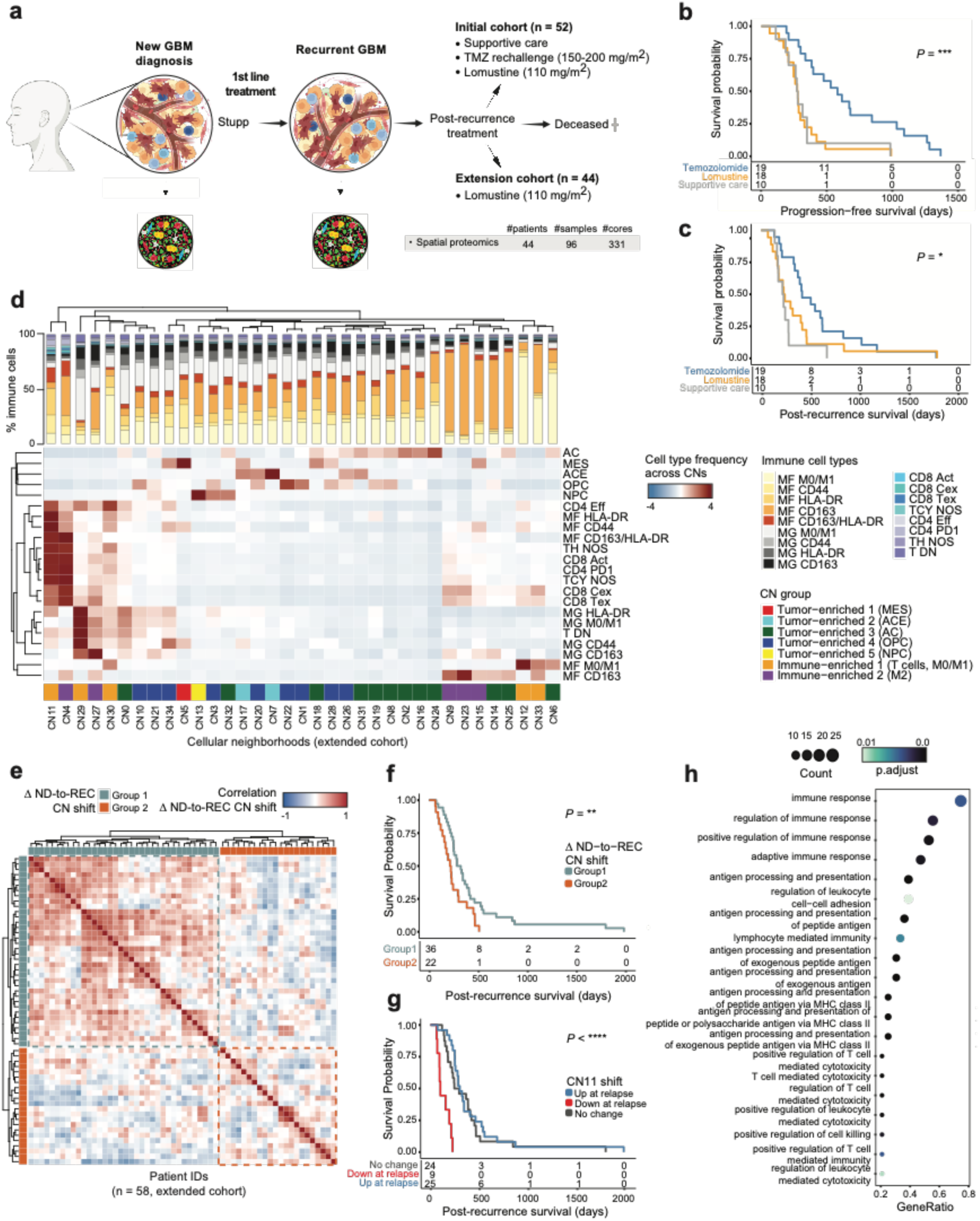
Impact of CNs ND-to-REC shift on post-recurrence treatment response and survival. **a** Overview of clinical course and post-recurrence treatment strategies in initial and extension cohort. See also Table S1b,c. TMZ: temozolomide. Image generated with Biorender.com. **b** Kaplan Meier plot showing differences in progression-free survival by post-recurrence treatment group in the initial cohort. Significance of differences was tested by Log-Rank test. ***, *P* ≤ 0.001. **c** Kaplan Meier plot showing differences in post-recurrence survival by post-recurrence treatment group in the initial cohort. Significance of differences was tested by Log-Rank test. *, *P* ≤ 0.05. **d** Normalized frequency of cell types across cellular neighborhoods (CNs) for lomustine-treated patients in the extended cohort (n = 58). The barplot annotation shows the percentage of the different immune cell types within each CN. AC, astrocyte-like; ACE, astrocyte-like (+EGFR expression); CD4eff, effector CD4+ T cells; CD4PD1, PD1+ CD4 T cells; CD8Act, activated CD8+ T cells; CD8Tex, terminally exhausted CD8+ T cells; CD8Cex, chronic exhausted CD8+ T cells; CNs, cell neighborhoods; DC, dendritic cells; MES, mesenchymal-like; MF, macrophages; MG, microglia; NOS, not otherwise specified; NPC, neural-progenitor-like; OPC: oligodendrocyte-progenitor-like; TCY, cytotoxic T cells; TDN, double-negative T cells; TH, helper T cells. **e** Patient stratification by Pearson correlation of the ND-to-REC shift in CN composition in the extended cohort (n = 58). CNs, cell neighborhoods; ND, newly diagnosed; REC, recurrent. **f** Kaplan Meier plot showing significant difference in post-recurrence survival between patient groups defined as of **e**. Significance of differences was tested by Log-Rank test. **, *P* ≤ 0.01. ND: newly diagnosed, REC: recurrent. **g** Kaplan Meier plot showing significant difference in post-recurrence survival between patient groups stratified by ND-to-REC shift in CN11 abundance. Significance of differences was tested by Log-Rank test. ****, *P* ≤ 0.0001. ND: newly diagnosed, REC: recurrent. **h** Dot plot showing GO terms overrepresented in CD163+/HLADR+ GAMs (spatial transcriptomics dataset). Dots are colored by the significance of the enrichment, while their size scales the count of the enriched genes. The x-axis highlights the gene ratio, *i.e.* the percentage of enriched target genes in each GO term.

As we observed that the evolution in the composition of cellular neighborhoods was associated with different survival patterns (Figure 3c,e-g), we investigated whether this characterization might also aid in better understanding the determinants of response to lomustine. We therefore extended our initial study cohort with 44 additional GBM patients, all of them treated with lomustine as part of their post-recurrence treatment scheme and for which longitudinal tumor material was available (Figure 5a, Supplementary Table 1b). In line with our initial observation, the progression-free survival of lomustine-treated patients in the extension cohort was significantly shorter [median survival 310 days (CI95% 287-426)] than that of patients receiving temozolomide rechallenge [571 days (CI95% 389-1070)] and similar to that of lomustine-treated patients in the initial cohort [271 days (CI95% 241-375)] (*P* = 7.80 x 10^-4^ by Log-Rank test), confirming the high medical need of this patient subset (Figure S6a).

Upon performing spatial proteomics analysis using a leaner 18-plex mIHC antibody panel that was designed to capture the most important cellular neighborhoods (Figure 5a, Table S14), we could recapitulate patterns of tissue organization and composition scored in the initial cohort (Figure 3a, Figure S6b), namely tumor-enriched CNs with relative dominance of specific tumor subtypes and varying abundance of immune cells as well as immune-enriched M0/M1 and M2-polarized CNs (Figure 5d). As before, we stratified patients based on their individual changes of cellular neighborhood composition at recurrence and identified two groups (Figure 5e) exhibiting different response to lomustine as second line treatment [post recurrence survival of 298 days (CI95% 245-398) vs 214 days (CI95% 163-295), *P* = 0.01, Log-Rank test], with a subset of patients (∼20%) surviving beyond 500 days (Figure 5f). When investigating whether particular changes in specific CNs might affect lomustine responsiveness, we found that a depletion of CN11 at recurrence was significantly associated with poor outcome as compared to no change or enrichment [median survival 99 days (CI95% 90-NA) vs 295 days (CI95% 251-454) for depletion and enrichment respectively, *P* < 1 x 10^-6^, Log-Rank test] (Figure 5g, S6c,d). This CN is primarily enriched in immune cells, particularly HLA-DR+, CD44+ and M0/M1 macrophages and, strikingly, CD163+/HLA-DR+ macrophages together with cytotoxic T cells as compared to other CNs (Figure 5d). Interestingly, we found that CD163+/HLA-DR+ glioblastoma-associated macrophages (GAMs) upregulated multiple pathways related to immune activation with ongoing antigen processing, presentation, and phagocytosis compared to the remaining GAMs (including HLA-DR+) (Figure 5h, Table S15, S16). This observation was further corroborated in single-cell expression data from Antunes *et al.*^13^, where a similar activation signature was observed in the macrophage subset (Figure S5e, Table S17, S18).

## Discussion

While several studies previously described how the cellular composition of GBM tumors can change from ND to REC at a population level^12,17,19,42^, most analyses have been constrained by small cohorts^43^, the lack of directly paired samples and lack of detailed patient and treatment characteristics^11,12^, the absence of spatial resolution^22^, or the integration of tumor cell biology with the microenvironment^9,22^. As a result, the precise trajectories of tumor evolution within individual patients and their alignment with clinical outcome have remained poorly resolved. By combining spatial single-cell profiling of paired samples with detailed clinical annotation, our study provides an integrated spatio-temporal map of GBM progression, uncovering specific, clinically relevant patterns of ecosystem remodeling. These insights enabled us to define a novel patient-stratification framework that correlates with multiple clinical parameters, including time to recurrence and post-recurrence responsiveness to lomustine in fast-recurring patients.

While recent (non-spatial) single-cell RNA-seq studies showed that transcriptional trajectories during GBM evolution are largely unpredictable and patient-specific^44,45^, we show that the evolution of cellular neighborhoods, i.e. spatially defined microenvironmental shifts that capture dynamic changes in tumor-immune-stromal interactions over time, aligns with clinical outcome and provide potentially actionable molecular insights. First, and in line with previous work, we found MES/AC-like tumor cells typically resided in a hypoxic environment^10^. Tumor cells in this niche expressed high levels of *CCL2*, a strong chemoattractant for macrophages that also induces a pro-tumoral/M2-like state, in addition to the induction of additional immunosuppressive mechanisms including high levels of TGFß, SPP1 and LGALS1^38–40^. In this process, TGFß and LGALS1 seem to play a dual role by enhancing pro-tumoral macrophage polarization, in combination with fueling mesenchymal transitions, proliferation and migration of tumor cells^38^. *SPP1*+ TAMs have previously also been shown to drive glioma progression^39^. For all three markers, interfering therapeutic strategies are in development although their clinical benefit remains unknown^38,40,46–48^. This multifaceted crosstalk underscores a self-sustaining ecosystem in which mesenchymal tumor cells and pro-tumoral macrophages co-evolve under hypoxic stress, promoting therapy resistance and disease progression.

Second, we also found that a transition towards an oligodendrocyte-progenitor-like (OPC-predominant) niche is associated with the most favorable clinical outcomes and the longest recurrence-free intervals, also aligning with *MGMT* promoter methylation. The latter is in line with previous work that suggested that *MGMT* methylation affects tumor composition^45^ and was most predictive for therapy response in high stemness tumors^49^. The OPC-like niche was characterized by an improved vascular structure, which is induced by tumor-derived *NPPC+* that stabilizes endothelial tight junctions and pericyte coverage in a PDGFB and CXCR3 dependent manner^36^, leading to a better oxygenated microenvironment and a balance between CCL3/4/IL10-driven immune modulation^41,50^ and MHCII-driven activity. Interestingly, systemic administration of recombinant NPPC in preclinical cancer models exerted therapeutic benefit by promoting vascular normalization and reducing hypoxia^34^, while NPPC was suggested to dampen inflammation by interfering with immune cell infiltration^34,51^.

Finally, we investigated whether the evolution in the composition of cellular neighborhoods would aid in better understanding the determinants of response to lomustine. The analysis of an extension cohort of another 44 patients that received post-recurrence lomustine treatment, confirmed that the retention of an immune-enriched, antigen-presenting niche at recurrence was predictive of lomustine sensitivity, whereas its loss was associated with rapid progression. Although lomustine is widely used as a standard therapy, there are still no prospectively validated biomarkers to identify the most eligible patients^5^. This suggests that tumors retaining activated myeloid populations appear better able to convert cytotoxic damage into better tumor control, whereas immune-niche loss fosters chemoresistance, suggesting that GBM treatment efficacy depends not only on tumor-intrinsic features (like MGMT methylation) but also on the spatial and ecosystem context at recurrence. Also, while current clinical practice rarely includes reoperation except in cases of symptomatic mass effect or trial enrolment, our results suggest that alterations in cellular ecosystem composition may serve as valid biomarkers, underscoring the importance of re-operation and tissue reassessment at recurrence^52^ .

Together, these findings provide a more granular understanding of how glioblastoma ecosystems evolve under therapeutic pressure and how these trajectories shape clinical outcome. By integrating longitudinal spatial single-cell profiling with clinical data, our study demonstrates that the direction of ecosystem evolution holds predictive value for patient prognosis and therapeutic responsiveness. Beyond refining patient stratification, these insights open avenues for targeted interventions that modulate the tumor microenvironment itself, moving towards adaptive, ecosystem-informed treatment strategies in glioblastoma.

## Materials and methods

### Patient cohorts

For this retrospective study, we reviewed the electronic medical records of GBM patients who were treated at the University Hospitals Leuven (Belgium), Ziekenhuis Oost-Limburg Genk (Belgium) and MUMC+ Maastricht (The Netherlands) between 12/2003 and 07/2021. Data collection and analysis were carried out in accordance with data protection guidelines. The coding system of the electronic medical database was used to select patients meeting the following criteria: histologically confirmed diagnosis of primary/de novo IDH^WT^ GBM and formalin-fixed paraffin-embedded (FFPE) tissue specimens available from at least two resections. Exclusion criteria were: (1) age younger than 18 years and (2) a lack of (good quality) tissue to perform further analyses (assessed by two expert neuropathologists - JB, RS). Ultimately, an initial discovery cohort of 52 patients and an expansion cohort of 44 patients with at least 2 resections during the course of their disease were included in the study (Figure 1a). In the expansion cohort, 49 patients were screened, from which 44 were analyzed and 40 patients had complete clinical information. After neurosurgical resection, SOC adjuvant treatment with radiotherapy and temozolomide (TMZ) was carried out. In the expansion cohort all patients received a lomustine-based regimen upon GBM recurrence. All patients were treated in routine clinical practice and were seen for regular clinical and radiological follow-up. The treating oncologist could change treatment approach concerning drug schedule, dose-reduction policy and timing of radiologic assessments in accordance with current local practice guidelines. Commonly used clinical features assessed for each patient are the age at diagnosis, sex, Karnofsky Performance Status (KPS), preoperative use of corticosteroids, location of the tumor, macroscopic and radiographic extent of resection (EOR), molecular pathological features of the tumor (*MGMT* promoter methylation, *EGFR* and *PDGFRA* amplification and *TP53* mutation, among others), adjuvant therapy, completion of Stupp protocol, time to recurrence, progression-free survival (PFS) and overall survival (OS) (Table S1). The study was approved by the medical ethics review board of all participating hospitals (*S59804, S61081, S62248, 18/0021R, 19/0021R, METC 16-4-022*).

### Spatial proteomics

#### MILAN Experimental Protocol

MILAN stands for ‘Multiple Iterative Labelling by Antibody Neodeposition’ and is co-developed by our lab^30,53,54^. It is a cyclic procedure where in each round, FFPE samples are (i) stained using an optimized immunofluorescent method, (ii) high resolution images are acquired of each fluorescent channel, and (iii) antibodies are stripped from the tissue according to a carefully developed method maximizing antigen preservation. Tissue slides are sequentially stained with a combination of three primary antibodies and 4’,6-diamidino-2-fenylindool (DAPI) to stain for cell nuclei. The technique makes use of fluorescently labelled secondary antibodies binding to the specific primary antibodies. After staining, a high-definition image is generated and then the antibodies are entirely removed using an SDS/ß-mercaptoethanol washing step to denature and inactivate the antibodies; after this, sometimes a bleaching step (Phosphate buffered saline/hydrogen peroxide/sodium hydroxide) is added to get rid of leftover signal. Afterwards, the next staining round with three new antibodies can be applied in a cyclic manner. By performing multiple rounds of stain-image-strip, multiplex panels of up to 80 antibodies can be applied to acquire single-cell data of a single FFPE tissue section^27,29,30,54,55^. For the list of antibodies used in this study, see Table S2-S3.

#### MILAN Computational Pipeline

Multiplex immunohistochemistry typically yields large data sets of multiple terabytes. Because tens to hundreds of parameters are measured simultaneously in thousands to millions of single cells across tens-of-thousands of images across numerous areas of a tissue, data analysis requires specific bioinformatics pipelines. To deal with the latter, a series of tailored algorithms enabling the processing of raw image files in a semi-automated way were developed and embedded in an integrated prototype software tool, named DISSCOvery (*Data integration of spatial single-cell omics by LISCO*) allowing image analysis, cell identification and digital reconstruction of the tissue by biology/pathology experts. First, several pre-processing steps are applied, where impurities and/or artifacts (e.g. dust, air bubbles, antibody precipitates, areas out of focus) are removed. Illumination corrections are applied, after which multiple small images are computationally stitched together to retrieve the single large image of the whole sample. The next step involves the exact superpositioning of images acquired from consecutive imaging rounds, commonly referred to as ‘image registration’. Background signal is removed to deal with autofluorescence (AF). Within each image, nuclei segmentation is then performed to ensure proper cell identification and cell type labelling. After the data analysis pipeline is completed, further analysis is executed in the program R and RStudio^56,57^.

#### Image Preprocessing

All acquired images underwent processing through the pipeline developed by our research group, as previously outlined in Antoranz et al.^58^. In summary, the preprocessing pipeline includes flat field correction, registration, segmentation, autofluorescence subtraction, and feature extraction. At each step, intermediate results were meticulously evaluated by manual inspection (MVM, PN, JB) for the identification of potential anomalies such as tissue folds, losses, external artifacts, and antibody aggregations. For the segmentation process, we employed Stardist^59^ using a pretrained model that underwent fine-tuning within our team. Nuclei were segmented, and topological as well as morphological features were extracted and incorporated into the dataframe for further analysis.

#### Blood Vessel Segmentation

Given the low staining intensity for CD31, a Convolutional Neural Network (CNN) with U-net architecture^60^ was trained to effectively segment these regions and generate corresponding masks (Figure S2). The training dataset comprised 20 carefully selected cores to ensure representation across the entire dataset. Following initial training, a final manual adjustment was conducted for specific cores where the generated masks did not meet the desired quality standards. This step aimed to enhance the accuracy and reliability of the segmentation results.

#### Autofluorescence Masks

To mitigate the risk of autofluorescent (AF) residues post-pipeline, potentially leading to false positivity for markers, additional masks were generated using the channel AF_FITC (fluorescein-5-isothiocyanate). The “autothresholdr” package in R, utilizing Renyi’s Entropy method, was employed for this purpose. The segmented regions underwent post-processing through morphological operations, including closing and filling holes (using the “EBImage” package in R). All resulting masks underwent manual evaluation (MVM, PN), with corrections made if necessary to ensure accuracy. To further enhance precision, the outer regions of tissues were incorporated into the aforementioned masks, excluding marginal areas where certain markers, such as GFAP, exhibited positivity. This comprehensive approach aimed to minimize the impact of autofluorescence and refine the segmentation results.

#### Cell Phenotyping

Following the exclusion of cells falling into the previously generated masks, marker expression values were *z*-scored per core and trimmed between -5 and 5 to mitigate the impact of potential outliers. Cell phenotyping (Figure S1) was performed using a hierarchical clustering approach: cells were first assigned to major lineages using *k*-means clustering, with the optimal number of clusters determined by Louvain community detection (PhenoGraph) to ensure stability^61,62^. Major cell compartments were defined using *SOX2, CD68, CD8, CD4, CD3,* and *TMEM119*, corresponding to tumor cells, macrophages, cytotoxic T cells (TCY), helper T cells (TH), and microglia. In a second step, each major lineage was further subdivided based on lineage-specific marker panels. Tumor cells were refined using *OLIG2, PDGFRA, CD44, VIM, NESTIN,* and *EGFR*; macrophages were subclustered using *CD163, HLADR,* and *CD44*; microglia was refined using the same marker panel. Cytotoxic T-cell subsets were defined by *LAG3, PD1,* and *GRB7*, while helper T-cell subsets were distinguished using *PD1* and *GRB7*.

#### Definition of Different Tumoral Areas

Through quadrant analysis, the cores were systematically categorized into four distinct regions based on the density of tumor cells. Cores were partitioned into squares measuring 70 by 70 micrometers, and the count of tumor cells as well as cells with no expressed markers (not otherwise specified (NOS) cells (=blanks)) in each square were used for quadrant allocation. Squares with >8 NOS cells were designated as “Necrotic.” For the remaining squares, the classification was as follows: squares with tumor cells greater than 8 were labeled as “High tumoral density,” those with 0 < N < 9 were categorized as “Low tumoral density,” and squares with N equal to 0 were classified as “Non-tumoral.” Thresholds and square size were determined through experimentation and visual inspection of the cores by neuropathologist experts (JB, RS), where the final results were validated by comparing digital reconstructions with hematoxylin and eosin (H&E) staining from consecutive slides (Figure 2a,b). This meticulous process aimed to accurately delineate and characterize different tumoral areas within the cores.

#### Definition of Neftel Regions

The minmax-normalized proportion of tumor subtypes and the total number of tumor cells within each square were considered for subsequent analysis. Using *k*-means clustering (k=30), the squares were grouped into clusters. These clusters were then meticulously annotated (MVM) and utilized to partition the cores into distinct tumoral subregions (oligodendrocyte-progenitor/OPC-like, neural-progenitor/NPC-like, mesenchymal/MES-like, astrocyte/AC-like and astrocyte with EGFR expression/ACE-like). This approach allowed for a nuanced characterization of tumor subtypes within the tissue samples, enhancing the understanding of the spatial distribution and composition of different tumor subtypes.

#### Cellular Neighborhoods (CNs)

To investigate spatial relationships between cell types, we applied the spatial_count function from the scimap Python package. This approach quantifies the local tissue microenvironment by enumerating neighboring cell types within a defined radial distance. Using X-Y coordinates, cell labels, and a 50-µm radius, CNs were identified by Euclidean distance, consistent with the quadrant analysis (√2 × 70/2 ≈ 49 µm). The resulting frequency matrix captured the distribution of neighboring cell types for each cell. For community analysis, we modelled these frequency profiles using latent Dirichlet allocation (LDA) to identify latent spatial motifs representing recurrent microenvironmental patterns across tissue regions. To further stratify communities, k-means clustering was applied. The number of clusters was selected based on a trade-off between the elbow method (inertia) and average silhouette width, ensuring an optimal balance between granularity and separation. Cells annotated as “NOS” were excluded from the analysis.

For the extension cohort, we assembled a lean panel of 18 markers that was assembled based on the community analysis in the discovery cohort, and that was still able to capture the key cellular phenotypes and their composing niches. Such lean panel would bear a higher translational value. Also, because the LOM expansion cohort contained fewer markers than the discovery dataset, cell types were grouped to achieve comparable phenotypic categories across cohorts. Specifically, macrophage classes *MF_CD11C* and *MF_M2* were merged with *MF_CD163*; *MF_CD206* was merged with *MF_M0/M1*; while *MF_CD163HLADR*, *MF_CD44,* and *MF_HLADR* were retained as distinct categories. For microglia, *MG_M2* was grouped with *MG_CD163*, and *MG_CD206* with *MG_M0/M1*, while *MG_CD44*, *MG_CD163*, *MG_M0/M1,* and *MG_HLADR* were preserved. Among T-cell subsets, *CD8Hsp* was merged with *CD8Tex* within the cytotoxic (TCY) compartment, and *CD4Hsp* and *Treg* were merged with *CD4PD1* within the helper (TH) compartment. *CD4Eomes* and *TH_NOS* were both mapped to *TH_NOS*. The dendritic cell (DC) class was excluded due to insufficient representation. After harmonizing cell type annotations, the initial and extension cohorts were combined, and CN scoring was performed jointly by spatial-LDA using the same parameters as in the initial analysis. Cluster selection followed the same optimization criteria (inertia and silhouette width), resulting in a final k = 35 for the integrated dataset. The alignment between CNs defined with either the original 38-plex (initial cohort) or limited 18-plex marker panel (extension cohort) was assessed by cosine similarity of CN composition (based on harmonized labels) on patients from the initial cohort (Figure S6b). Grouping of CNs in the extension cohort was assigned by matching each CN with the most similar CN from the initial analysis, followed by label transferring.

### Spatial transcriptomics

#### Nanostring CosMx Experimental Protocol

Nanostring CosMx SMI^63^ is an integrated system with mature cyclic fluorescent in situ hybridization (FISH) chemistry, high-resolution imaging readout, interactive data analysis and visualization software. A tissue micro-array (TMA) was constructed including a subset of 14 carefully selected GBM tissue pieces of 2mm-diameter, encompassing 7 patients with both a resection at diagnosis and upon recurrence. A balanced subset of tissue samples was analyzed to preserve tumor materials. Fresh-cut sections of 5 μm thickness were mounted on a VWR Superfrost Plus Micro Slide (Cat#: 48311-703). After sectioning, the sections were air-dried at room temperature overnight. Afterwards, slides were permeabilized and fixed, after which hybridization of RNA specific probes and antibodies was applied, followed by flow cell assembly. After the hybridization and imaging of a reporter set, each round is followed by UV cleavement and washing of the fluorescent dyes before the next reporter set is applied. In total, a panel of 1008 genes was measured at the spatial, single-cell level (Table S6). After measuring the RNA transcripts, a count table is generated for the field of views that have been analyzed.

#### Nanostring CosMx Analysis

Single-cell spatial transcriptomic analysis was performed using the Seurat toolkit (v5.1.0). As initial quality control, cells expressing <10 or >500 genes were filtered out. Preprocessing steps included normalization and variance stabilization using the SCTransform method, hypervariable features selection (n = 250), dimensional reduction by principal component analysis and Uniform Manifold Approximation and Projection (UMAP) on 40 principal components and 30 nearest-neighbors. Initial cell type profiling was performed upon nearest-neighbor graph construction and clustering (Louvain algorithm) to annotate endothelial, immune and tumor cells (*i.e.* level 1 annotation, Figure S5a,c, Table S8). This annotation was further refined by subclustering of immune and tumor cells and validated by checking the expression of canonical markers (*i.e.* level 2 annotation, Figure S5b,d, Table S9). Marker genes for differential expression analysis were computed by the FindAllMarkers function using the Wilcoxon Rank-Sum test, a minimum fraction of at least 0.25 cells in either of contrasting populations. Multiple test adjustment of p-values was performed using Bonferroni correction; only significant genes (p < 0.05) with at least 0.25 positive log fold change were retained for subsequent analyses. Gene Ontology (GO) enrichment analysis of differentially expressed genes was performed using clusterProfiler (v4.8.1).

To integrate transcriptional and proteomic data from consecutive slides, manual registration was performed to align the CosMx fields of view with the corresponding MILAN regions. The alignment was guided by visual inspection of tissue architecture and the spatial organization of cell phenotypes. The level of concordance between the two modalities was formally assessed by computing, for each sample, the cosine similarity over cell type proportions by aligned tiles (using harmonized labels) and using the mean of optimally matched similarities as summary metric (median = 0.94) (Table S7). Based on level 2 cell type annotation, cellular neighborhoods (CNs) were independently computed on spatial transcriptomics data using the same approach described for spatial proteomics. For each CN, dominance of endothelial, immune and tumor cells was established based on the proportion of level 1 cell types and Pearson correlation analysis of level 2 cell type composition (Figure S5e,f). Cytokine polarization of tumor-enriched CNs was scored based on clustering of scaled gene expression of *CCL2* vs *NPPC* (Figure 4c).

### Integrated Spatial Proteomics and Transcriptomics by RNAscope™ and seqIF™ profiling

#### Spatial Multiomics on COMET™ by Lunaphore Technologies

Automated hyperplex multiome immunofluorescence staining and imaging were obtained on the COMET™ platform by combining the RNAscope™ HiPlex Pro Assay and sequential immunofluorescence (seqIF™)^64^ assays for the detection of 24 protein markers and 12 RNAscope™ probes (Table S10). The OME-TIFF output contains nuclear staining (DAPI), four autofluorescence channels, and the 14 protein markers and 12 RNAscope™ channels to be used for downstream analyses.

#### Sample Pre-processing

FFPE sections of 3 µm were captured on positively charged slides, baked overnight at 56°C, and underwent dewaxing using the Leica ST5010 Autostainer XL (Leica Biosystems) where slides transitioned through a graded xylene and ethanol series. Further pre-processing for antigen retrieval was performed on PT model (Epridia) with HIER Buffer H (AR03, Lunaphore Techologies) for 60 min at 99°C. Subsequently, slides were washed and stored with Multistaining Buffer (BU06, Lunaphore Technologies) until later use.

#### HiPlex Pro and SeqIF™ sample processing

The 12-plex RNA plus 24-plex protein protocol was generated using the COMET™ Control Software where reagents were loaded onto the instrument to perform the multiome protocol by combining the automated RNAscope™ HiPlex Pro Assay (322075, Advanced Cell Diagnostics) to co-detect RNA with SeqIF™. Samples were pretreated in the RNAscope™ HiPlex PretreatPro buffer for 30 min at 40°C, followed by the hybridization of the RNAscope™ HiPlex probe cocktail for 2 hours at 40°C. After washing the samples with Multistaining Buffer (BU06, Lunaphore Technologies) at 37°C, the probe signal was amplified by consecutive hybridization of RNAscope™ HiPlex Pro AMP1, AMP2, AMP3 oligos for 30min at 37°C. Amplified signals were detected after incubation with RNAscope™ HiPlex Pro Fluoro 4-channel detection probes for 15 min at 37°C. Images were detected in channels FITC, TRITC, Cy5, Cy7, followed by fluorophore cleavage using RNAscope™ HiPlex Pro Cleaving Reagent diluted in 4X Saline Sodium Citrate for 15min at 27°C. Detection and cleavage steps were repeated for 3 cycles until all target probes were detected. After completion of the RNAscope™ HiPlex assay, the seqIF™ protocol was started.

List of used RNAscope™ probes and primary antibodies is enclosed in Table S10. Secondary antibodies were used as a mix of two species-complementary antibodies supplemented with DAPI (Table S10). All reagents were diluted using Multistaining Buffer (BU06, Lunaphore Technologies). For each cycle, the following exposure times were set up: DAPI 25 ms, TRITC 250 ms, Cy5 400 ms. Elution steps of 2 minutes were performed with Elution Buffer (BU07, Lunaphore Technologies) at 37°C. Signal quenching using the quenching step lasted for 30 sec using the Quenching Buffer (BU08, Lunaphore Technologies). Resulting output of the multiomics protocol reveals a multi-layer OME-TIFF file where the post-elution imaging outputs of each cycle are stitched and aligned.

### Genetics

The genetic background of the patient samples was defined with either TSO500 next-generation sequencing (NGS) panel (22/52 samples – MUMC+, NL) or whole-exome sequencing (WES) (19/52 samples - Novogene, UK). TSO500 NGS was performed as described in Heuvelings et al. ^65^. WES has been performed on Illumina Novaseq 6000^66^. After sample preparation, a quality control step was applied, followed by exome capture, library preparation, library QC, sequencing and data quality control before bioinformatics analysis. We also performed a methylation-specific PCR to define the *MGMT* promoter methylation status for the entire cohort, both on the ND and the REC samples, given the known predictive impact for response on alkylating chemotherapy. The Oncoprint graph (Figure 1c) summarizes the most relevant genetic background and clinical information of the paired glioblastoma patient cohort.

### Statistics

All statistical analyses were performed in R (version 4.0.5) using RStudio (version 2021.0.9). No statistical methods were used to pre-determine sample sizes. Comparisons between ND and REC groups were carried out using the non-parametric Wilcoxon signed-rank test for paired samples. For survival analyses, patients who had not experienced death at the time of analysis or were lost to follow-up were censored at the date of last progression. Linear correlations were assessed using the Pearson correlation coefficient (*r*). Delta values (Δ) denote the difference in proportion of features of interest (either cell types or cellular neighborhoods) between ND and REC samples for each individual patient. To account for multiple testing, *P*-values were adjusted using the Bonferroni method unless otherwise specified. Statistical significance was defined as adjusted *P* < 0.05.

### Survival analysis

Survival analysis was performed using the R packages survival (v3.8.3) and survminer (v0.5.0). Differences in survival outcomes by different grouping variables were visualized with the Kaplan Meier estimator method; p-values for statistical significance were estimated by Log-Rank test. To assess differences for continuous variables (Figure S6c), optimal split was determined using the surv_cutpoint function.

To identify which CNs (based on their ND-to-REC shift) were best predictors of post-relapse lomustine response, we performed the Log-Rank test independently for each CNs (using time from 2^nd^ resection to death as survival time). CNs with p-values < 0.1 were selected to fit a LASSO-penalized Cox regression model using the glmnet R package (v4.1.8). Model stability was assessed over 100 iterations; at each iteration, performance was evaluated via cross-validated concordance index (C-index), reported as mean ± standard deviation. CNs consistently identified as predictor in at least 80% of iterations were then encoded as categorical variables to represent the direction of the shift: “no change” (defined as within 0.5 × median absolute deviation around zero), “enrichment” (above threshold), or “depletion” (below threshold).

## Data availability

The source data supporting findings in this study have been deposited at KU Leuven RDR: https://doi.org/doi:10.48804/2JCUJS. Raw primary imaging data can be obtained from the authors directly upon reasonable request.

## Code availability

Code is available https://gitlab.kuleuven.be/explore/projects

## Acknowledgements

MVM is supported by an FWO PhD fellowship fundamental research (11L0822N & 11L0824N). AA is supported by an FWO Postdoctoral fellowship. FDS, BD, FB, and SDV are supported by KOTK grants (KOTK/2018/11509/2 and KOTK/2019/11892/1). This research was supported by the Horizon Europe initiative (grant 101073386 - www.gliomatch.eu), funded by the European Union. Views and opinions expressed are however those of the author(s) only and do not necessarily reflect those of the European Union or the European Health and Digital Executive Agency (HaDEA). Neither the European Union nor the granting authority can be held responsible for them. Other funding agencies supporting this work include Research Foundation Flanders (FWO; including grant #G0B3722N, S001221N, I005920N and G0I1118N), and the KULeuven Bijzonder onderzoeksfonds (CELSA/20/022). We would like to thank all members of LISCO – KULeuven Institute for Single Cell Omics, in particular the platform for multiplexed immunohistochemistry - https://lisco.kuleuven.be/MILAN. We wish to thank medical students Bram Janssen, Bram Swinnen, Berne Florus, Fleur Vercammen and Eline Beyen for their contributions. We wish to thank all the patients, family members and staff from all the units that participated in the study.

## Author contributions

<u>Conceptualization</u>: MVM, JB, SDV, PC, MV, RS, EJS, AH, FDS

<u>Data curation:</u> MVM, PN, JB, CC, DL, BC, GS, MV, SB, TM, SZ, EG, AA, JP, AH

<u>Formal analysis:</u> MVM, PN, JB, CC, DL, BC, CBGB, GS, DP, JT, AA, JP

Funding acquisition: MVM, EJS, AH, FDS

<u>Investigation:</u> MVM, PN, JB, CC, DL, BC, CBGB, BD, YDV, MV, DP, JT, JM, YVH, AC, MD, FB, RS, AA, JP, EJS

<u>Methodology:</u> MVM, PN, JB, DL, BC, GS, FB, RS, AA, JP, EJS, AH, FDS

Project administration: MVM, JB, AH, FDS

<u>Resources:</u> JB, DE, AZH, FW, TD, PVE, MG, FM, KH, SDV, PC, MB, MV, RS, EJS, AH, FDS

<u>Software:</u> MVM, PN, GS, IS, AA, JP

<u>Supervision:</u> JB, PC, EJS, AH, FDS

<u>Validation:</u> MVM, PN, JB, DL, BD, YDV, MV, DP, JM, YVH, AC, MD, FB, RS, AA, JP, EJS

<u>Visualization:</u> MVM, PN, JB, CC, DL, BC, GS, FB, AA, JP

<u>Writing – original draft:</u> MVM, AH, FDS

<u>Writing – review and editing:</u> all co-authors

