## Supplementary information for "Longitudinal tumor ecosystem mapping defines glioblastoma treatment trajectories"

### Supplementary figures

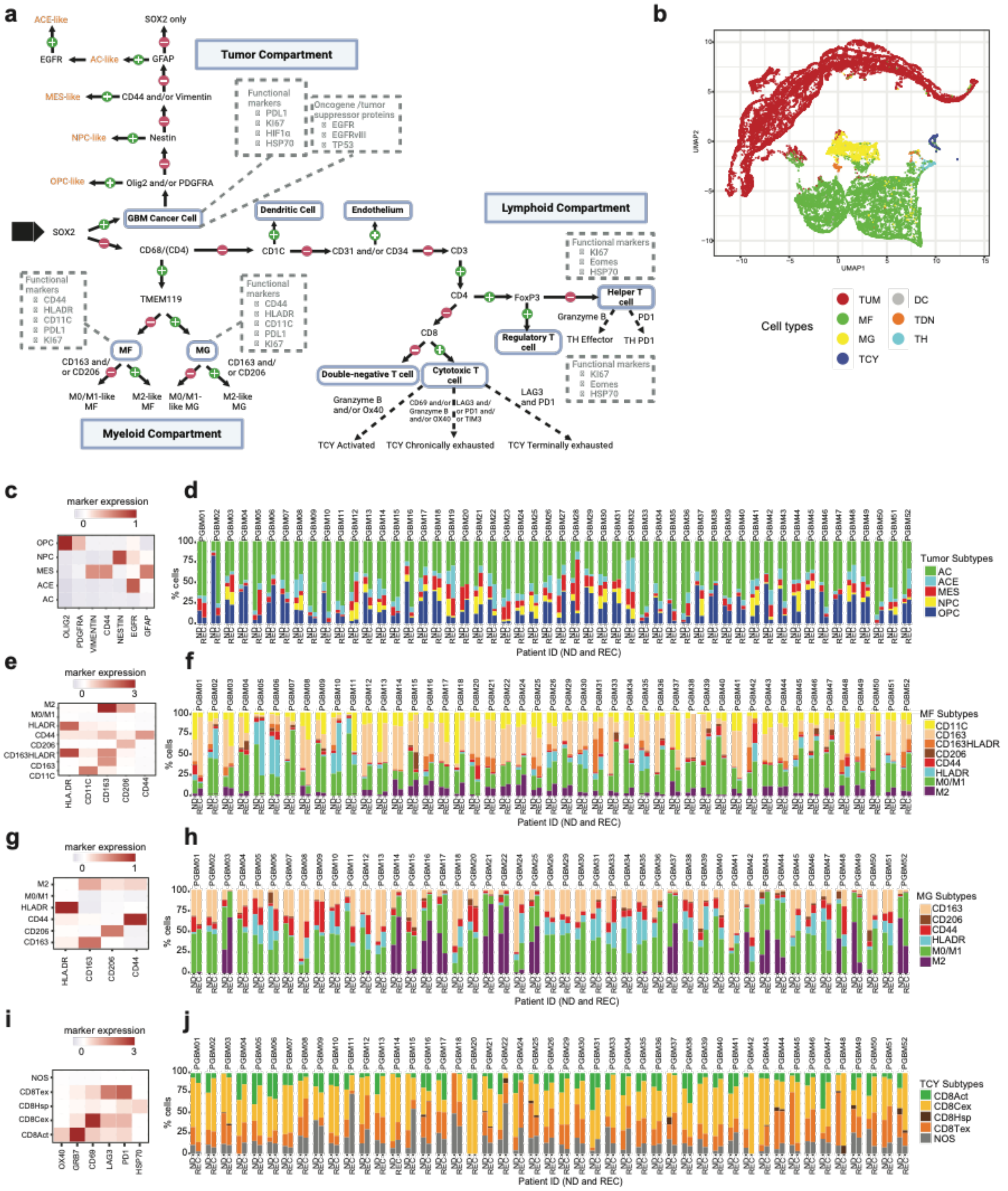


**Figure S1: Overview of patient cohort and experimental approach**

**a** Flowchart of the markers stained in MILAN analysis, showing the different annotated cell types. First, main cell types were clustered using a hierarchical approach (CD3, CD4, CD8, Sox2, CD68, TMEM119) and within each main cell type we clustered the cells with phenotypical/functional markers to define subtypes. We took into account the positivity/negativity of the markers in the clusters to label them as a certain cell type. Image generated with Biorender.com.

**b** UMAP of the main cell types. DC: dendritic cells, MF: macrophages, MG: microglia, TCY: cytotoxic T cells, TDN: double negative T cells, TH: helper T cells, TUM: tumor cells.
**c** Heatmap defining different SOX2+ tumor cell subtypes based on marker expression. Kmeans clustering is used for clustering analysis. The AC-like tumoral subtype was defined by exclusion since GFAP is a very stable marker, abundantly present in nearly all the analyzed glioblastoma samples. OPC: oligodendrocyte-progenitor-like, NPC: neural-progenitor-like, MES: mesenchymal-like, ACE: astrocyte-like (+EGFR expression), AC: astrocyte-like.
**d** Overview of the proportion of tumoral subtypes across the 52 patients, both in the ND and REC samples. Different cores from the same timepoint have been pooled. Patients are ordered from shortest overall survival (PGBM01) until the longest overall survival (PGBM52). AC: astrocyte-like, ACE: astrocyte-like (+EGFR expression), MES: mesenchymal-like, NPC: neural-progenitor-like, OPC: oligodendrocyte-progenitor-like.
**e** Heatmap defining different CD68+/TMEM119- macrophage subtypes based on marker expression. Kmeans clustering is used for clustering analysis. M2-polarized macrophages are positive for both CD163 and CD206. M0/M1 macrophages are defined by exclusion if no other (M2) markers were expressed.
**f** Overview of the proportion of macrophage subtypes across the 52 patients, both in the newly diagnosed and recurrent samples. Different cores from the same timepoint have been pooled. Patients are ordered from shortest overall survival (PGBM01) until the longest overall survival (PGBM52). MF: macrophages.
**g** Heatmap defining different CD68+/TMEM119+ microglia subtypes based on marker expression. Kmeans clustering is used for clustering analysis. M2-polarized MG are positive for both CD163 and CD206. M0/M1 MG are defined by exclusion if no other (M2) markers were expressed.
**h** Overview of the proportion of microglia subtypes across the 52 patients, both in the ND and REC samples. Different cores from the same timepoint have been pooled. Patients are ordered from shortest overall survival (PGBM01) until the longest overall survival (PGBM52). MG: microglia.
**i** Heatmap defining different CD3+/CD8+ T cell subtypes based on marker expression. Kmeans clustering is used for clustering analysis. NOS: not otherwise specified, CD8Tex: terminally exhausted T cells, CD8Hsp: Hsp70+ expressing T cells, CD8Cex: chronically exhausted T cells, CD8Act: activated T cells.
**j** Overview of the proportion of cytotoxic T cell subtypes across the 52 patients, both in the newly diagnosed and recurrent samples. Different cores from the same timepoint have been pooled. Patients are ordered from shortest overall survival (PGBM01) until the longest overall survival (PGBM52). TCY: cytotoxic T cells.

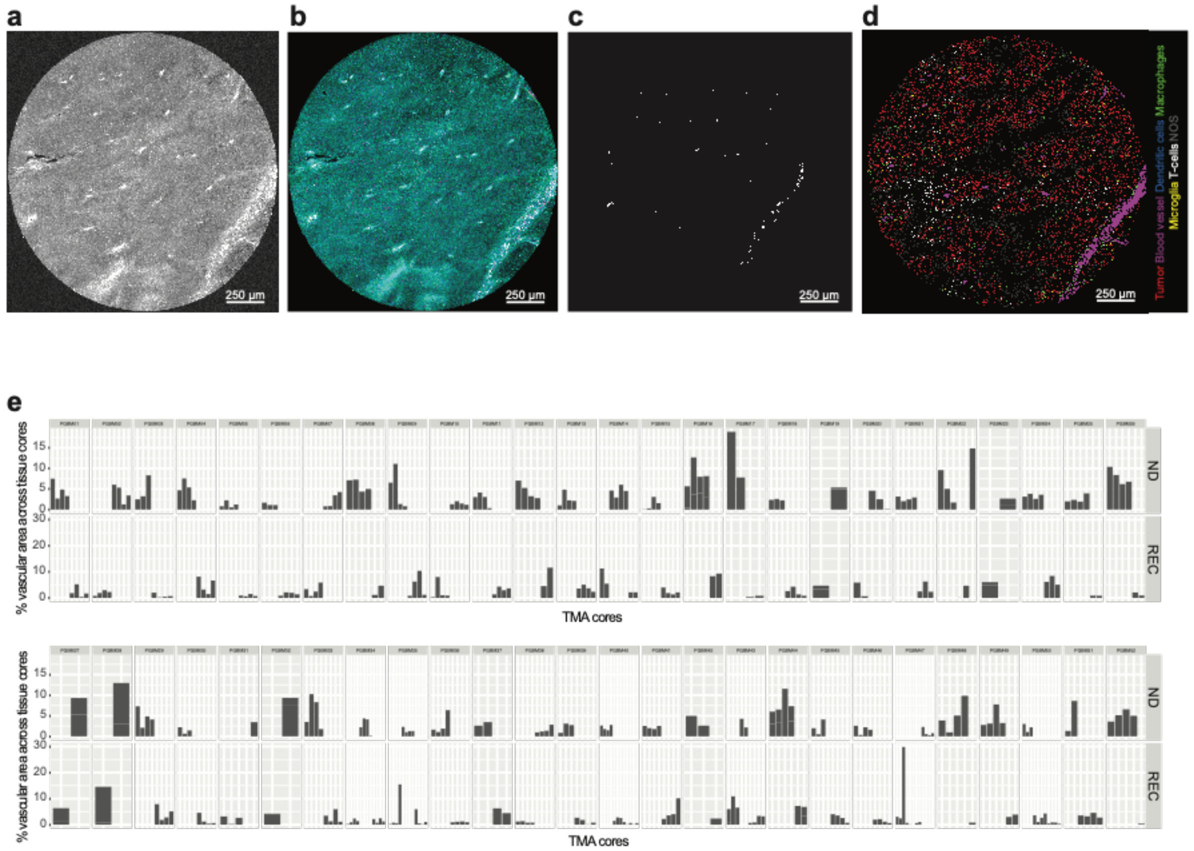


**Figure S2: Identification and quantification of blood vessels from spatial proteomics**

**a-e** Main steps for blood vessel identification from spatial proteomics images on one example tissue core. For details on the analysis pipeline, see also *Materials and methods*.

**a** Autofluorescent image. Red blood cells show a higher level of autofluorescence compared to other tissue structures.

**b** CD31 protein staining (in purple).

**c** Blood vessel mask.

**d** Digital reconstruction depicting the main cell types.

**e** Percentage of vascular area across all tissue cores analyzed for the 52 patients. ND newly diagnosed; REC, recurrent.

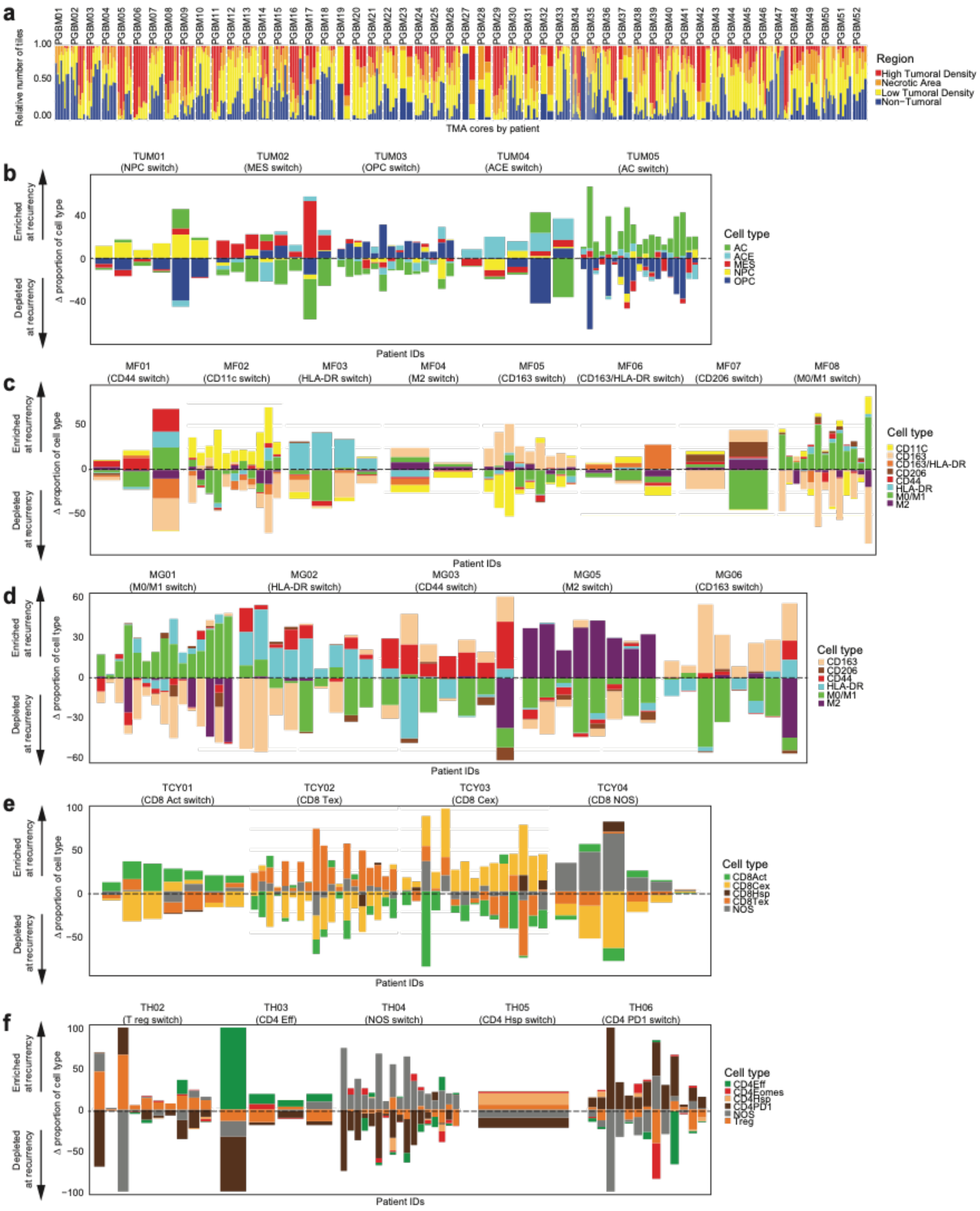


**Figure S3: Patterns of ND-to-REC evolution in cell type composition and correlation to progression-free survival**

**a** Proportion of different GBM tissue regions for each single tissue core for each patient. Regions were classified based on the number of tumor cells in a square micro-region of 70 x 70 micron: 'High tumoral density' (tumor cells > 8), 'Low tumoral density' (0 < tumor cells ≤ 8), 'Necrotic area' (NOS cells > 8) and 'Non-tumoral area' (no tumor cells).

**b-f** Patterns of evolution in cell type composition of the low-tumor density regions between newly diagnosed (ND) and recurrent (REC) tumor samples for each individual GBM patient. The delta value (Δ, on the x-axis) shows the ND-to-REC difference in the proportion of each cell type, allowing to group patients with similar shifts into different evolutionary patterns. (**b**) Tumor subsets; TUM, tumor;

AC, astrocyte-like; ACE, astrocyte-like (+EGFR expression); MES, mesenchymal-like; NPC, neural-progenitor-like; OPC: oligodendrocyte-progenitor-like. (**c**) Macrophage subsets; MF, macrophages. (**d**) Microglia subsets; MG, microglia. (**e**) Cytotoxic T cells subsets; TCY, cytotoxic T cells; CD8Act, activated CD8+ T cells; CD8Tex, terminally exhausted CD8+ T cells; CD8Cex, chronic exhausted CD8+ T cells; NOS, not otherwise specified. (**f**) T helper cells subsets; TH, helper T cells; CD4Eff, effector CD4+ T cells; Treg, regulatory T cells; NOS, not otherwise specified.

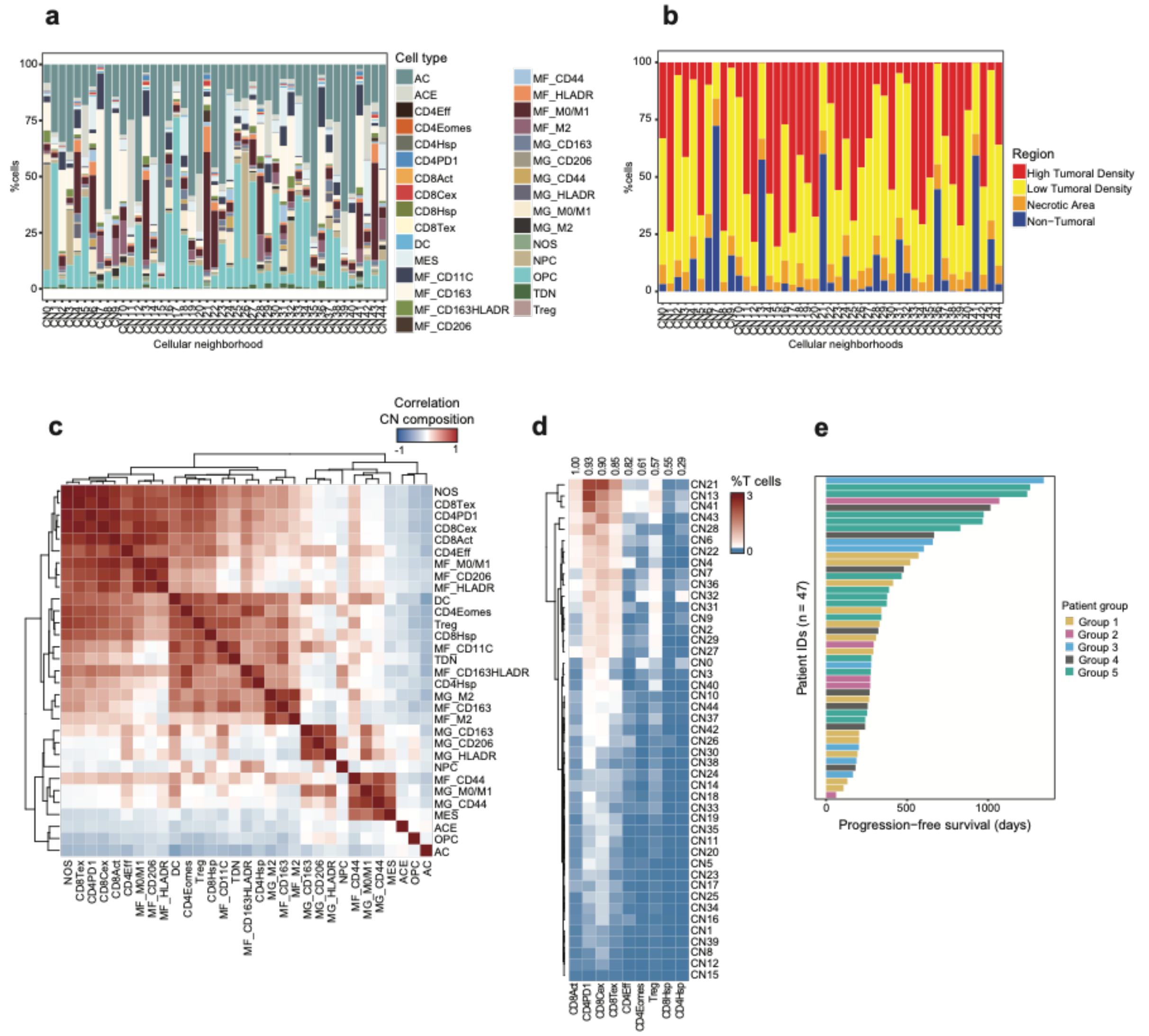


**Figure S4: Patterns of ND-to-REC evolution in cellular neighborhood composition and correlation to progression-free survival**

**a** Proportion of cell types across the 45 cellular neighborhoods (CNs). AC, astrocyte-like; ACE, astrocyte-like (+EGFR expression); CD4eff, effector CD4+ T cells; CD4PD1, PD1+ CD4 T cells; CD8Act, activated CD8+ T cells; CD8Tex, terminally exhausted CD8+ T cells; CD8Cex, chronic exhausted CD8+ T cells; DC, dendritic cells; MES, mesenchymal-like; MF, macrophages; MG, microglia; NOS, not otherwise specified; NPC, neural-progenitor-like; OPC: oligodendrocyte-progenitor-like; TCY, cytotoxic T cells; TDN, double-negative T cells; TH, helper T cells; TUM, tumor.

**b** Proportion of high/low-density tumoral, non-tumoral and necrotic areas across the 45 cellular neighborhoods (CNs).

**c** Pearson correlation and clustering analysis on the cellular composition of the cellular neighborhoods, highlighting the colocalization of specific cell types. AC, astrocyte-like; ACE, astrocyte-like (+EGFR expression); CD4eff, effector CD4+ T cells; CD4PD1, PD1+ CD4 T cells; CD8Act, activated CD8+ T cells; CD8Tex, terminally exhausted CD8+ T cells; CD8Cex, chronic exhausted CD8+ T cells; DC, dendritic cells; MES, mesenchymal-like; MF, macrophages; MG, microglia; NOS, not otherwise specified; NPC, neural-progenitor-like; OPC: oligodendrocyte-progenitor-like; TCY, cytotoxic T cells; TDN, double-negative T cells; TH, helper T cells; Treg, regulatory T cells.

**d** Proportion of T cell subtypes across the cellular neighborhoods (CNs) where CD8Act cells are present. Pearson correlation was computed between proportion of CD8Act cells and each of other T cell subtypes to highlight co-localization patterns (shown in the top annotation). CD4eff, effector CD4+ T cells; CD4PD1, PD1+ CD4 T cells; CD8Act, activated CD8+ T cells; CD8Tex, terminally exhausted CD8+ T cells; CD8Cex, chronic exhausted CD8+ T cells; Treg, regulatory T cells.

**f** Progression-free survival of GBM patients based on subgroups with similar ND-to-REC shift in CN composition (as defined in Figure 3c). CN, cellular neighborhoods; ND, newly diagnosed; REC, recurrent.

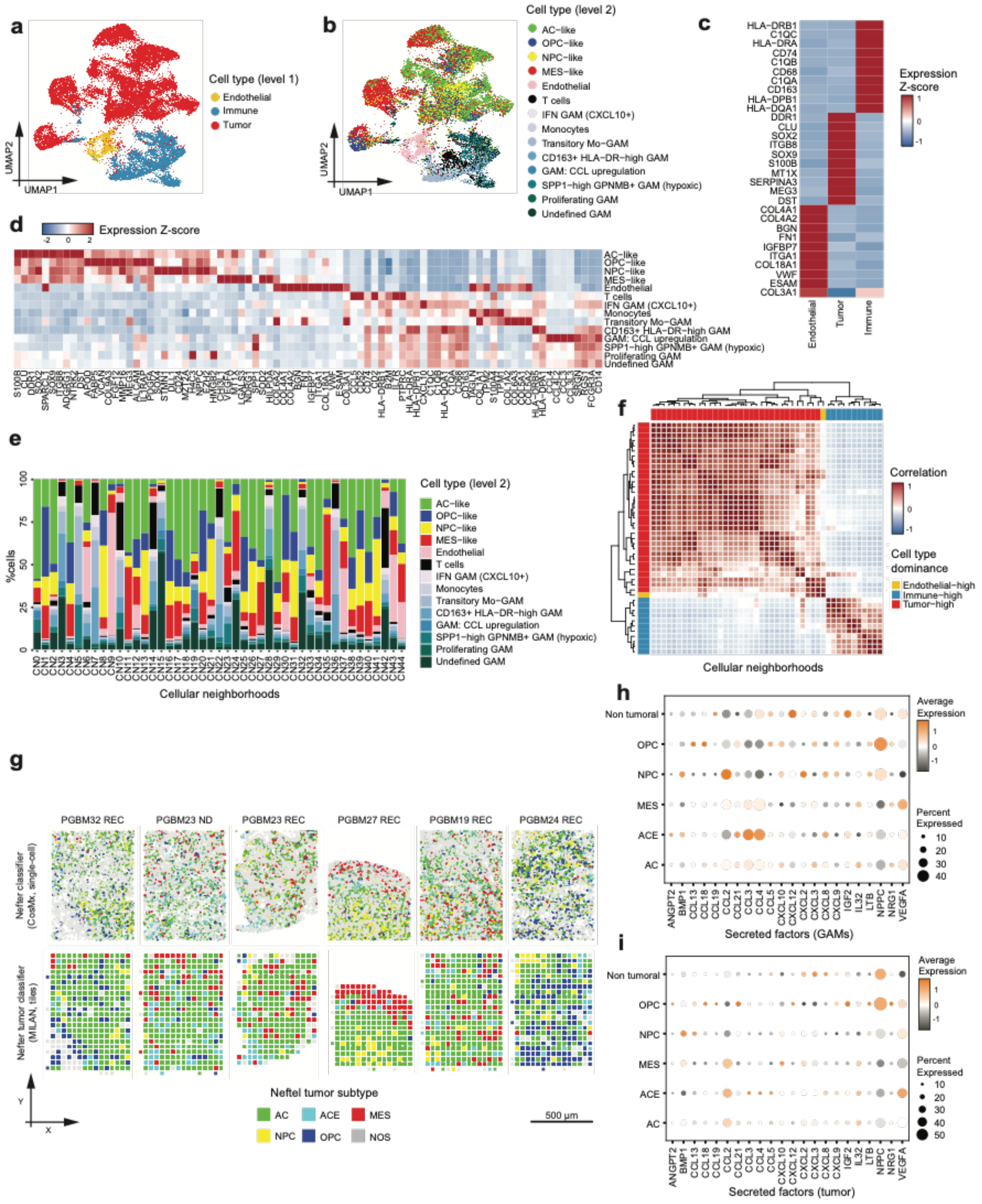

**Figure S5: Mapping niche-specific cytokine networks**

**a** UMAP plot showing level 1 cell type annotations from spatial transcriptomics data.

**b** UMAP plot showing level 2 cell type annotations from spatial transcriptomics data. AC, astrocyte-like; DC, dendritic cells; GAMs, Glioblastoma-associated macrophages; IFN, interferon; MES, mesenchymal-like; Mo, monocytes; NPC, neural-progenitor-like; OPC: oligodendrocyte-progenitor-like.

**c** Top 10 markers of level 1 cell types from spatial transcriptomics data.

**d** Top 10 markers of level 2 cell types from spatial transcriptomics data. AC, astrocyte-like; DC, dendritic cells; GAMs, Glioblastoma-associated macrophages; IFN, interferon; MES, mesenchymal-like; Mo, monocytes; NPC, neural-progenitor-like; OPC: oligodendrocyte-progenitor-like.

**e** Proportion of level 2 cell types by CNs from spatial transcriptomics data. AC, astrocyte-like; CN, cellular neighborhood; DC, dendritic cells; GAMs, Glioblastoma-associated macrophages; IFN, interferon; MES, mesenchymal-like; Mo, monocytes; NPC, neural-progenitor-like; OPC: oligodendrocyte-progenitor-like.
**f** Pearson correlation of cellular neighborhoods (CNs) based on cell type composition as defined from spatial transcriptomics data. Clustering denotes CNs enriched in tumor, endothelial or immune cells (Tumor-high, Endothelial-high and Immune-high dominance respectively) as indicated in the top annotation bar.
**g** Integrating spatial transcriptomics (top row) and spatial proteomics for which local tumor subtype enrichments are highlighted (bottom row) in consecutive tissue slides. For the tumoral compartment, the Neftel classifier has been applied as described in *Methods*. A subset of 6 highly correlating cores has been used for representation. AC: astrocyte-like, ACE: astrocyte-like (+EGFR expression), GAMs: Glioblastoma-associated macrophages, MES: mesenchymal-like, NOS: not otherwise specified, NPC: neural-progenitor-like, OPC: oligodendrocyte-progenitor-like. ND: newly diagnosed, REC: recurrent.
**h-i** Expression of various secreted factors in the integrated spatial transcriptomic data set in tumor cells **(h)** and GAMs **(i)** across the various tumor subtype areas, as well as non tumor areas. AC: astrocyte-like, ACE: astrocyte-like (+EGFR expression), MES: mesenchymal-like, NPC: neural-progenitor-like, OPC: oligodendrocyte-progenitor-like.

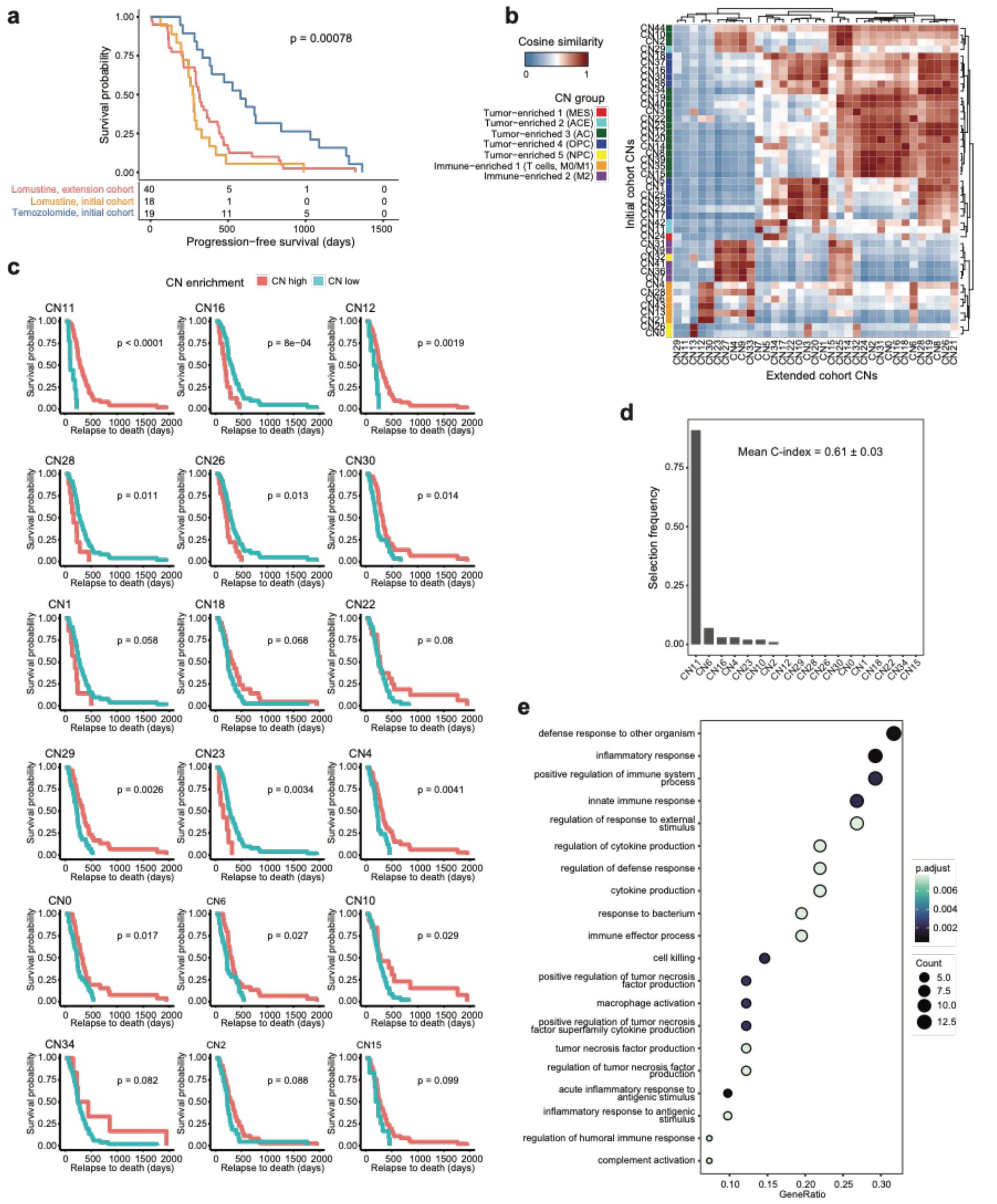

**Figure S6: Impact of CNs ND-to-REC shift on post-recurrence treatment response and survival**

**a** Kaplan Meier plot of progression-free survival for GBM patients grouped by post-recurrence treatment strategy (either temozolomide rechallenge or lomustine) and study cohort (initial or extended cohort). Significance of differences was tested by Log-Rank test.

**b** Heatmap showing alignment between CNs defined for the initial and extended cohort respectively, assessed by cosine similarity of CN composition (based on harmonized labels). CNs from the initial cohort are grouped as in Figure 3a. CNs: cellular neighborhoods.

**c** Kaplan Meier plots showing probability of post-recurrence survival based on optimal split in the ND-to-REC shift of each CNs. The panel shows CNs associated with difference in survival ($P < 0.1$ as estimated by Log-Rank test). CNs: cellular neighborhoods.

**d** Lasso-penalized Cox regression on CNs selected from **c** to identify best predictors of response to lomustine rescue in fast-recurring GBM patients. The barplot shows the frequency of selection of each tested CN across 100 iterations. Concordance index (C-index) is reported as mean ± standard deviation. CNs: cellular neighborhoods.
**e** Dot plot showing GO terms overrepresented in CD163+/HLADR+ GAMs (single-cell RNA dataset from Antunes *et al.*[13]). Dots are colored by the significance of the enrichment, while their size scales the count of the enriched genes. The x-axis highlights the gene ratio, *i.e.* the percentage of enriched target genes in each GO term.

### Supplementary table legends

**Table S1a: Patient characteristics (initial cohort; full)**
Extended patient demographics, treatments and outcomes metadata.

**Table S1b: Patient characteristics (initial cohort; summary)**
Summary of patient demographics, treatments and outcomes. mOS: median overall survival, mPFS: median progression-free survival, MGMT: O-6-methylguanine-DNA methyltransferase, ND: newly diagnosed, REC: recurrent.

**Table S1c: Patient characteristics (extension cohort; summary)**
Summary of patient demographics, treatments and outcomes. mOS: median overall survival, mPFS: median progression-free survival, MGMT: O-6-methylguanine-DNA methyltransferase, ND: newly diagnosed, REC: recurrent.

**Table S2: MILAN antibody list (broad; initial cohort)**
Overview of antibodies used in the MILAN multiplex IHC analysis including clone, reference, host, concentration and vender company.

**Table S3: MILAN antibody list (small; initial cohort)**
Overview of antibodies used in the MILAN multiplex IHC analysis including clone, reference, host, concentration and vender company. The smaller panel has been applied on a consecutive slide on the subset of 14 tissue cores that have been analyzed with spatial transcriptomics.

**Table S4: Pairwise Log-Rank test for CN-based patient subgroups**
Pairwise Log-Rank test for differences in progression-free survival between GBM patient subgroups with similar ND-to-REC shift in CN composition.

**Table S5: Fisher's exact test for genetic features by CN-based patient subgroups**
Fisher's Exact Test for associations between genetic features and GBM patient subgroups with similar ND-to-REC shift in CN composition.

**Table S6: Spatial single-cell transcriptomics gene panel (CosMx)**
Full panel of 1008 genes measured with Nanostring CosMx SMI.

**Table S7: Mean cosine similarities between aligned CosMx and MILAN datasets**
For each tissue samples profiled with spatial transcriptomics and proteomics on consecutive sections, mean of optimally matched cosine similarities over cell type proportions by aligned tiles (using harmonized labels).

**Table S8: Marker genes of level 1 cell type annotation (CosMx)**
Marker genes for differential gene expression analysis of different level 1 cell type annotations were computed by the FindAllMarkers function using the Wilcoxon Rank-Sum test, a minimum fraction of at least 0.25 cells in either of contrasting populations, and multiple test adjustment of p-values by Bonferroni correction.

**Table S9: Marker genes of level 2 cell type annotation (CosMx)**
Marker genes for differential gene expression analysis of different level 2 cell type annotations were computed by the FindAllMarkers function using the Wilcoxon Rank-Sum test, a minimum fraction of at least 0.25 cells in either of contrasting populations, and multiple test adjustment of p-values by Bonferroni correction.

**Table S10: List of used RNAscope™ probes and primary antibodies**
List of 12 RNAscope™ probes and primary antibodies for 24 protein markers used to combine the RNAscope™ HiPlex Pro Assay and sequential immunofluorescence (seqIF™)[64] assays.

**Table S11: Marker genes of CCL2-high vs NPPC-high, tumor enriched cellular neighborhoods for endothelial cells, glioblastoma-associated macrophages and T cells (CosMx)**
Marker genes of each tumor-enriched cellular neighborhood (labelled as CCL2-high or NPPC-high) were computed separately for endothelial cells, glioblastoma-associated macrophages and T cells by the FindAllMarkers function using the Wilcoxon Rank-Sum test, a minimum fraction of at least 0.25 cells in either of contrasting populations, and multiple test adjustment of p-values by Bonferroni correction.

**Table S12: Post-recurrence therapy**
Overview of the systemic treatments that were given at disease progression after resection at disease recurrence.

**Table S13: Fisher's exact test for *MGMT* promoter methylation status by post-recurrence therapy**
Fisher's Exact Test for associations between *MGMT* promoter methylation status and post-recurrence treatment strategy.

**Table S14: MILAN antibody list (extension cohort)**
Overview of antibodies used in the MILAN multiplex IHC analysis including clone, reference, host, concentration and vender company.

**Table S15: Marker genes of CD163+/HLA-DR+ macrophages vs other glioblastoma-associated macrophages (CosMx)**
Marker genes of CD163+/HLA-DR+ macrophages vs other glioblastoma-associated were computed by the FindAllMarkers function using the Wilcoxon Rank-Sum test, a minimum fraction of at least 0.25 cells in either of contrasting populations, and multiple test adjustment of p-values by Bonferroni correction.

**Table S16: Gene Ontology (GO) enrichment analysis of CD163+/HLA-DR+ macrophages marker genes**
List of enriched pathways resulting from over representation analysis of CD163+/HLA-DR+ macrophages marker genes.

**Table S17: Marker genes of CD163+/HLA-DR+ macrophages vs other glioblastoma-associated macrophages (Antunes *et al.*)**
Marker genes of CD163+/HLA-DR+ macrophages vs other glioblastoma-associated extracted from single-cell RNA sequencing data (Antunes *et al.).*

**Table S18: Gene Ontology (GO) enrichment analysis of CD163+/HLA-DR+ macrophages marker genes (Antunes *et al.*)**
List of enriched pathways resulting from over representation analysis of CD163+/HLA-DR+ macrophages marker genes from Antunes *et al.*

### Supplementary Notes

##### Note S1 – Clinical description of the analysed patient cohorts

##### Initial cohort (n=52 patients)

In this work, we have assembled a multicentric, retrospective cohort of 52 GBM patients provided with tumor tissue (FFPE) from both new diagnosis (ND) and recurrence (REC) across three different clinical centers (UZ Leuven and ZOL Genk from Belgium, and Maastricht UMC+ from Netherlands). Of all included patients, detailed longitudinal clinical and radiological information in addition to treatment characteristics were collected from the institutions' databases (see methods) starting from initial diagnosis to the moment of death. Most patients (50/52) received standard of care treatment as per the international guidelines with the Stupp regimen (60 Gy radiotherapy in 2 Gy daily fractions for 6 weeks with concurrent TMZ 75 mg/m$^2$ 7 days per week, followed by 6 cycles adjuvant TMZ 150-200 mg/m$^2$, 5 days during each 28-day cycle)[1] (Figure 1a). Only two patients in the outlined cohort were treated with radiation therapy alone. Recurrence was defined as radiological reappearance of tumor after total resection or tumor progression in case of subtotal resection. The longitudinal design and homogenous first-line treatment schedule allow to reliably correlate changes in tissue architecture with treatment outcomes in this study cohort.

The demographic/clinical characteristics and outcomes of all included patients are outlined in Figure S1 + Table S1a,b and are representative of real-life GBM patients. Overall, the mean age at diagnosis was 57 years (range 26-80 years), with a male predominance (65%). The most prevalent tumor location was in the frontal lobe (50%), followed by the temporal lobe (37%). Expectedly, patients often received preoperative corticosteroids (90% of ND resections, 72% of REC resections). *MGMT* promoter hypermethylation was present in 22 patients at initial diagnosis and in 17 patients at recurrence, and, in line with previous reports[67,68], *MGMT* promoter methylation became more prevalent with longer OS. The median PFS of our patient cohort was 13.1 months (10.2-16.0 months 95% CI) with a median OS of 28.1 months (23.4-32.8 months 95% CI). The presence of *MGMT* promotor hypermethylation at diagnosis results in a PFS of 14.7 months (9.6-19.7 months 95% CI) and OS of 34.8 months (27.1-42.5 months 95% CI); which is significantly longer compared to the *MGMT* unmethylated counterparts (PFS 11.9 months, OS 23.4 months). When comparing ND and REC samples, in seven patients a switch in *MGMT* promoter methylation was found, where six of them loose their initial hypermethylation at recurrency.

It is important to note that this cohort focused on a subgroup of GBM patients that were able to endure multiple resections (typically 20-40% of patients, dependent on the clinical center[69–72] , explaining the somewhat increased survival time compared to historical survival rates typically described in GBM).[1] Following a more detailed analysis of treatment schemes, patients were further divided in subgroups consisting of those that completed the entire treatment protocol (6 cycles, n = 32), those with only partial completion of the adjuvant TMZ cycles (1-5 cycles, n = 18), or those without adjuvant TMZ treatment (n= 2). Causes for early discontinuation of TMZ treatment were the occurrence of significant hematological toxicity or disease progression during adjuvant treatment. While the median OS of patients with a total and subtotal extent of resection (EOR) at diagnosis was comparable (28.0 months vs 30.1 months respectively), the amount of applied adjuvant cycles of TMZ and the OS of the patients did correlate: patients who received all 6 out of 6 adjuvant TMZ cycles had a significantly longer OS than patients who received 1-5 adjuvant TMZ cycles or radiotherapy alone (33.2 vs 18.7 months, p=0.0006). No significant differences in OS were observed between patients that did not receive TMZ therapy (concurrent and adjuvant) and patients who received partial (1 to 5 cycles of) adjuvant TMZ in our study group (p=0.096). At the same time, patients who were able to receive all 6 cycles of TMZ tended to exhibit a higher KPS at diagnosis. Combinations of the following treatment options were given at times of progression: chemotherapy (rechallenge of TMZ or other salvage chemotherapy schedules containing lomustine), re-resection, re-irradiation, immunotherapy, angiogenesis inhibitors (e.g. Bevacizumab) or study medication (Table S1a,b).

To further characterize this cohort, genomic profiling (see methods) was done on DNA extracted from FFPE materials (n = 41 patients; for 11 patients DNA quality was insufficient). Overall, we identified classical GBM-related genomic aberrations, including $EGFR^{AMP/MUT}$, $PTEN^{MUT}$, $TP53^{MUT}$, $PDGFRA^{AMP}$, $MDM2/4^{AMP}$, $CDK4^{AMP}$, and $BRAF^{MUT}$ according to previously described frequencies[28], and these could either be retained, appear, or disappear between ND and REC in a patient-specific pattern (Figure 1c). No enrichment for specific genomic aberrations was found in the aforementioned subgroups.

**Extension cohort (n=44 patients)**

Additionally, we have assembled a multicentric, retrospective cohort of 44 GBM patients provided with tumor tissue (FFPE) from both new diagnosis (ND) and recurrence (REC) across the three collaborating clinical centers (UZ Leuven and ZOL Genk from Belgium, and Maastricht UMC+ from Netherlands). Of all included patients, detailed longitudinal clinical and radiological information in addition to treatment characteristics were collected from the institutions' databases (see methods) starting from initial diagnosis to the moment of death. Most patients (41/44) received first-line standard-of-care treatment according to the international guidelines with the Stupp regimen (Figure 1a). Only 3 patients in this lomustin expansion cohort were treated with radiation therapy alone in the first line setting. Recurrence was defined as radiological reappearance of tumor after total resection or tumor progression in case of subtotal resection. All included patients receive a (partial) reresection at tumor recurrence; which was followed by a lomustin containing chemotherapy treatment schedule: lomustin monotherapy (30%) or PCV (70%) (lomustin (110 mg/m2, d1), procarbazin (60 mg/m2/d, d8-d22) and vincristin (1,4 mg/m2, d8, d29)). The demographic-and clinical characteristics and also the outcomes of all included patients in the lomustine expansion cohort are outlined in Table S1b and are representative of real-life GBM patients. Overall, the mean age at diagnosis was 58 years (range 30-84 years), with a male predominance (75%). The most prevalent tumor location was in the temporal lobe (45%), followed by the frontal lobe (32%). The median overall survival of this IDHwt GBM cohort was 23 months (ranging from 9-93 months). The median progression free survival was 11 months (ranging from 1 upto 43 months). Only in 10 patients (23%) *MGMT* promoter methylation status at initial diagnosis was defined. No additional molecular analysis of this expansion cohort was available.
